# The genetic architecture and evolutionary consequences of the human pelvic form

**DOI:** 10.1101/2024.05.02.592256

**Authors:** Liaoyi Xu, Eucharist Kun, Devansh Pandey, Joyce Y. Wang, Marianne F. Brasil, Tarjinder Singh, Vagheesh M. Narasimhan

## Abstract

Human pelvic shape has undergone significant evolutionary change since the divergence from the chimpanzee lineage. This transformation, involving the reduction of the pelvic canal size to support bipedal locomotion, is thought to give rise to the obstetrical dilemma, a hypothesis highlighting the mismatch between the large brain size of infants and the narrowed birth canal in females. Empirical evidence for this classic hypothesis has been equivocal, largely due to a lack of sample size and appropriate types of data. To elucidate the genetic underpinnings of pelvic morphology, we applied a deep learning model to 31,115 dual-energy X-ray absorptiometry (DXA) from the UK Biobank, extracting a set of seven pelvic proportion (PP) phenotypes, including measures of the birth canal. All PPs were found to be highly heritable (∼25-40%) and a genome-wide association study of these traits identified 179 independent loci. Unlike other skeletal proportions including long bone lengths, the subpubic angle associated with the birth canal exhibits a genetic correlation between sexes significantly less than 1, in line with sex-specific reproductive function. PPs were also left-right asymmetric but not heritable and instead associated with handedness. We conducted phenotypic and genetic association analyses to link PPs to 3 facets of the dilemma: locomotion, pelvic floor function and childbirth. Larger birth canal phenotypes were associated with reduced walking pace, decreased risk of back pain, and increased risk of hip osteoarthritis - phenotypes linked to locomotor efficiency. We also observed that a narrower birth canal width was associated with a reduced risk of pelvic floor disorders. When examining childbirth-related outcomes, narrower birth canal phenotypes were associated with increased risk of emergency cesarean sections and obstructed labor due to insufficient dilation, but not obstructed labor due to positioning of the fetus. Finally, we examined whether the dilemma might have been alleviated through evolution. We found no association between any PPs and gestational duration, contrary to the initial prediction by Washburn in 1960. However, we found that the birth weight of the child, a proxy for skull and brain size, was genetically correlated with birth canal width but not with other PPs. Collectively, our study offers fresh insight on a 60-year-old debate in human evolutionary studies. Our results support the idea that the obstetrical dilemma has played a central role in the co-evolution of the human brain and pelvis, while also highlighting the potential role of associated factors such as pelvic floor health.

## Introduction

The human skeleton has undergone significant morphological change associated with the transition to bipedalism. Some of the most significant changes occurred in the pelvis, resulting in a superoinferiorly short and mediolaterally flaring pelvis relative to the modern great apes (*1*, *2*). These features are believed to have emerged early in hominin evolution and the alteration in pelvic anatomy allowed for the positioning of the upper body above the lower limb joints and facilitated the maintenance of an upright posture (*3*). While debate continues about the details of gait mechanics in fossil hominins (*1*) it is clear that the modern human pelvis is adapted to habitual bipedality, and undergoes a specific pelvic motion during walking that is thought to reduce energetic costs associated with bipedal locomotion (*4*).

The suite of adaptations for bipedality includes a reduction of the bi-acetabular distance, minimizing pelvic rotation during bipedal movement and consequently enhancing efficiency (*5*). This narrowing of the bi-acetabular distance results in a narrower birth canal, and is thought to stand in direct opposition to the birthing of children with significantly larger brains than our evolutionary predecessors (e.g., (*6*–*14*)). In the 1960s, this functional and evolutionary conflict was coined the “obstetrical dilemma” by Washburn (*13*). In the six decades since then, the obstetrical dilemma has been a source of intense debate, and different studies have attempted to examine the validity of the hypothesis through empirical data (*6*, *14*–*17*). One area of contention centers on the relationship between pelvic shape and walking efficiency or walking speed. Some studies have found there is an association between the two (*7*, *18*), while others have not (*19*–*22*). Another point of debate revolves around whether differential birth canal proportions are associated with obstruction during delivery (*7*–*14*, *17*, *23*–*26*). Recently, there has been growing appreciation for the concept of a multifactorial pelvis, which proposes that the role of pelvic width reduction is not just to enable bipedal locomotion, but also to reduce the risk of pelvic floor disorders. Pelvic width reduction improves the pelvic floor’s ability to support the fetus and the inner organs, and to prevent incontinence (*7*, *27*, *28*).

In addition to debates about the association between pelvic morphology and locomotion, childbirth, and pelvic floor function, it has been suggested that in modern humans the obstetrical dilemma has been alleviated through evolution. Washburn’s initial hypothesis proposed that relative to the other great apes, humans experience a shorter gestation period. This enables human infants to be born relatively earlier in development than their primate counterparts, consequently limiting the extent of brain growth before birth and ensures that the newborn can successfully traverse the birth canal during delivery. However, this hypothesis has been challenged and updated in recent years, as human gestational length and newborn size have been found to align with or exceed expectations for primates of our size, similar to the other great apes (*14*, *29*–*31*) (see (*6*, *7*, *32*) for alternate usages and historical perspectives on the term “obstetrical dilemma”).

While different aspects of the dilemma have been tackled over the past few decades, these previous studies suffer from several shortcomings. One issue with many studies – particularly those involving clinical outcomes – is that measurements of pelvic dimensions were collected externally (*19*, *22*), which may not adequately reflect the skeletal constraints imposed, particularly with respect to the birth canal. Another issue is that some earlier studies lack complete information about individual lifetime health records and are unable to distinguish between fine-grained but important details such as elective and emergency C-sections. However, the major challenge contributing to the ongoing debate is the limited sample size in many of these studies, which often only have data on a few hundred individuals (sample sizes and references of previous papers are reported in **Table S1**. In addition, data obtained for each study is often only capable of addressing one facet of the dilemma, as datasets examining childbirth outcomes and pelvic morphology often do not include data about pelvic floor function or walking speed/efficiency for the same individuals.

Finally, the underlying basis of skeletal evolution in the pelvis is genetic. While functional genomic datasets examining gene expression through development as well as comparative gene expression between the great apes and humans for the pelvis have yielded valuable insights (*33*–*35*) study of the direct association between pelvic trait variation and genetics has not yet been carried out. Thus, the genetic basis of pelvic morphology underlying variation in humans or indeed any other vertebrate is largely unknown, precluding analysis of natural selection on pelvic phenotypes directly at the genomic level.

Here, we applied methods in computer vision to derive a comprehensive set of seven skeletal measurements of the human pelvis from full-body dual-energy X-ray absorptiometry (DXA) images at biobank scale. We performed genome-wide scans on these seven phenotypes to identify loci associated with variation in pelvic proportions (PPs). Using summary statistics from these image-derived phenotypes (IDPs), we linked human PPs through phenotypic and genetic correlation with other biobank phenotypes, with an emphasis on locomotor, pelvic floor and childbirth-related outcomes.

## Results

### A deep learning approach to measure pelvic morphology

To study the genetic basis of the human pelvis, we jointly analyzed DXA and genetic data from 42,284 individuals in the UK Biobank (UKB). Individuals from this dataset are between 40 and 80 years old and reflect adult skeletal morphology. We report baseline information about our analyzed cohort in **Table S2**. Using a previously published deep learning-based image quality control (QC) pipeline (*36*), we retained only DXA images for the full body which included the entire pelvis, and removed images which contained image artifacts, atypical aspect ratios, and other abnormalities, retaining 39,469 images of high quality. These images were then uniformly cropped and padded to focus on the pelvis for subsequent analysis (**Methods**: *A deep learning model to identify pelvic landmarks on DXA scans*).

After performing image QC, we manually annotated 17 landmarks on 293 randomly selected pelvic images (see **Fig. S2**) to train our model. To assess the accuracy of our manual annotations, we re-annotated 20 images from the initial set of 293 and refined this annotation through model-in-the-loop labeling (**Methods**: *Image quality control*, **Fig. 1B** and **1C**). Our deep learning model was based on a High-Resolution Network (HRNet) architecture chosen because it maintains a high-resolution representation throughout the model which improves the performance of landmarking for this task on benchmarking tasks. These methods were robustly applied to a similar task of identifying joints on the overall skeleton (*36*, *37*) (**Methods**: *Image quality control*).

**Fig. 1.**
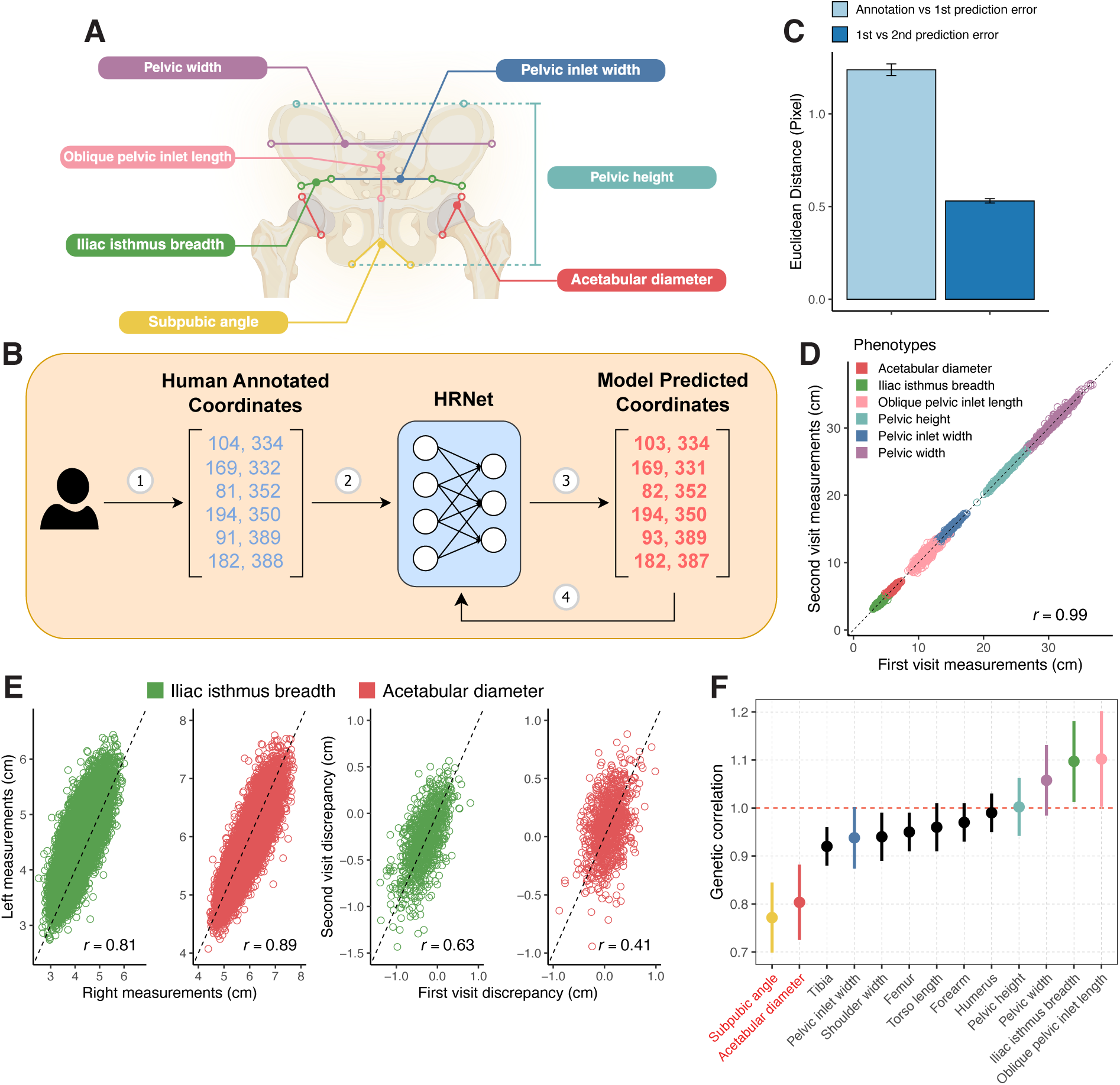
Deep learning-based image quantification and validation. (**A**) Deep learning-based image landmark estimation using the HRNet architecture is shown. During this process, 293 training images manually annotated with specific landmarks were used to train the model, which to perform automatic annotation of landmarks on the rest of images in the dataset from which pelvic measurements were calculated. (**B**) Model in the loop training data workflow. The coordinates from the 293 training images initially annotated by humans were used as a training set to train a model that was then redeployed on the training data. This helped to remove variation present in human labelling of the images and refined the training data itself. (**C**) Model in the loop training reduces annotation variability. Light blue bar indicates the average Euclidean distances between human annotated landmarks and the model’s first prediction on 58 validation set images. The dark blue bar indicates the average Euclidean distance between first and second model prediction on 58 validation set images. (**D**) Correlation of lengths measured from the first and second imaging visits for the same individual. (**E**) The two panels on the left side show the correlation between the left- and right-side measurements of the iliac isthmus breadth and acetabular diameter. The two panels on the right side illustrate the correlation of the left-right discrepancy in the iliac isthmus breadth and acetabular diameter between the first and second imaging visit. (**F**) Genetic correlation between female and male pelvic phenotypes and other skeletal traits including tibia, femur, torso length, forearm, and humerus. The error bars show 1 standard error. Heritability greater than 1 is due to small sample size. The two traits shown in red on the x-axis are the only ones that are significantly different from one.

### Validation of human pelvic phenotype estimates

After training and validating the deep-learning model on the 297 manually annotated images, we applied this model to predict the 17 landmarks on the rest of the 39,469 full-body DXA images. We then calculated the pixel Euclidean distances between pairs of landmark coordinates to ascertain six length phenotypes: pelvic width, pelvic inlet width, oblique pelvic inlet width, iliac isthmus breadth, pelvic height and acetabular diameter, and one angle phenotype: subpubic angle (**Fig. 1A**). To standardize images with varying aspect ratios, we rescaled pixels into centimeters for each image resolution. This was achieved by regressing the pixel height against the standing height in centimeters, as measured in the UK Biobank assessments (**Methods**: *Image standardization*, **Fig. S11**). For all seven pelvic measurements, we excluded individuals exceeding four standard deviations from the mean (**Methods**: *Removal of image outliers*, **Fig. S11**).

Following outlier removal, we validated the accuracy of our measurements on the remaining samples in two ways. First, we calculated the average error between labels in the validation data and model performance: average error was 2 pixels across all 17 landmarks. Second, we analyzed 935 individuals with repeat imaging visits at least two years apart. The correlation of all pelvic length phenotypes between the first and second imaging visits was greater than 0.99 (**Fig. 1D**). This indicates that the phenotype estimations via our deep learning model are both accurate and highly replicable.

### Human pelvic asymmetry is associated with handedness, and is not heritable

Next, we examined the correlation between measurements on the left and the right side of the pelvis. The two phenotypes with measures on each side were iliac isthmus breath and acetabular diameter. The left-right correlation for iliac isthmus breadth and acetabular diameter were 0.809, and 0.894 respectively (**Fig. 1E**). The average difference between the measurements in the iliac isthmus breadth between the left and right sides was 0.287 cm (*p* < 2 × 10^-16^, 95% confidence interval (CI) = 0.294 to 0.280), and for acetabular diameter, it was 0.101 cm (*p* < 2 × 10^-16^, 95% CI = 0.093 to 0.108). Though these differences were small, we found that they were replicable - left and right discrepancies in individuals across two imaging visits had Pearson correlations of 0.633 and 0.407 for iliac isthmus breadth and acetabular diameter respectively (**Fig. 1E**). This suggests that we can capture a measure of pelvic asymmetry beyond measurement error. On estimating the heritability of this trait using GCTA (*38*) we found that it was consistent with 0 (hg^2^ for acetabular diameter discrepancy = 0.0131, SE = 0.0149, hg^2^ for iliac isthmus breadth discrepancy = 0.0275, SE = 0.0158). However, we observed a significant association between pelvic asymmetry and handedness - another trait that is also not significantly heritable (left-handed hg^2^ = 0.0104, right-handed hg^2^ = 0.0096 in 150,000 individuals). The genetic correlation between acetabular diameter discrepancy and left-handedness is -0.39, and with right-handedness, it is 0.34. Similarly, the genetic correlation between iliac isthmus breadth discrepancy and left-handedness is 0.15, and with right-handedness, it is 0.11. In addition, we regressed the left and right pelvic phenotype ratio against handedness while controlling for age and sex. Right-handed individuals tended to have larger right acetabular diameters than left-handed individuals (regression *p* = 8.31 × 10^-6^) and larger left iliac isthmus breadth than left-handed individuals (regression *p* = 0.0665). This suggests that left-right pelvic asymmetry might be driven by left- or right-side dominance which is itself not heritable, but affect movement patterns and consequently skeletal development.

### Sexual dimorphism in the genetic basis of PPs

The human pelvis plays a critical role in childbirth and is one of the most dimorphic skeletal elements between males and females (*39*–*41*). Given the distinct functionalities between male and female pelvis, we examined whether the genetic basis of our seven pelvic phenotypes differed between males and females. To do so, we carried out genetic correlation analysis between a GWAS carried out in males versus females. Functionally similar pelvic phenotypes, such as pelvic height, exhibit similar genetic architectures between males and females, with a genetic correlation of 1.03. In contrast, birth canal-related phenotypes like the subpubic angle showed genetic correlations significantly divergent from 1. This difference in genetic correlation is in striking contrast to virtually all other skeletal traits previously examined such as arm, leg, torso, and shoulder dimensions. These other traits all showed genetic correlations not significantly different from 1 in the same cohort (**Fig. 1F**), suggesting that sex-specific reproductive requirements of the human birth canal are driving genetic differences between sexes for these PP traits.

### GWAS of human PPs

We performed GWASs using imputed genotype data in the UKB to identify variants associated with each pelvic phenotype. We applied standard variant and sample QC and focused our analyses on 31,115 individuals of “white British ancestry,” as defined by the UKB genetic assessment, and 7.4 million common biallelic single-nucleotide polymorphisms (SNPs) with minor allele frequency >1%. We used BOLT-LMM (*42*) to regress variants on each skeletal measure using a linear mixed-model association framework. We included height as a covariate to directly adjust for differences in body size between individuals and focus on skeletal proportions instead of overall length. We also adjusted for body size differences in two other ways: dividing each phenotype by height to generate a skeletal proportion, and including a leave one-chromosome-out polygenic risk score (PRS) for height as a covariate in the GWAS (*43*). GWAS effect sizes using either height as a covariate or height combined with the one-chromosome-out PRS as a covariate were highly correlated (Pearson correlation = 0.99) (**Methods**: *Adjusting for height correlation in GWAS by adding height as covariate*, **Fig. S13**). For downstream analyses, we focused on the results from the GWAS that included height as a covariate. Notably, we show that the effect sizes estimated for our PP phenotypes were uncorrelated with effect sizes from height (**Methods**: *Adjusting for height correlation in GWAS by adding height as covariate*, **Fig. S14**, average Pearson correlation across all phenotypes = 5.67 × 10^−5^, standard deviation = 0.0097), suggesting that PPs and height are distinct traits.

After generating summary statistics for each skeletal measure, we estimated SNP heritability using LD Score regression (LDSC) (*44*) and GCTA-REML (*38*). All traits were highly heritable, with SNP heritability between 25% and 40% for LDSC and between 17% and 50% for GCTA-REML (**Methods**: *GWAS and Heritability analysis*, **Fig. 2B**, **Fig. S15**). Across the six pelvic phenotypes adjusted by height (pelvic width, pelvic height, iliac isthmus breath, acetabular diameter, pelvic inlet width, oblique pelvic inlet length) and subpubic angle, we identified 339 loci at *p <* 5 × 10^−8^ and 241 loci at *p* < 7.14 × 10^−9^ (Bonferroni correction for seven traits). Of these loci, 179 are independently significant at *p* < 5 × 10^−8^ (linkage disequilibrium (*r*^2^) < 0.1) across all seven phenotypes (119 after Bonferroni correction for seven traits at *p* < 7.14 × 10^−9^) (**Fig. *2*A**).

**Fig. 2.**
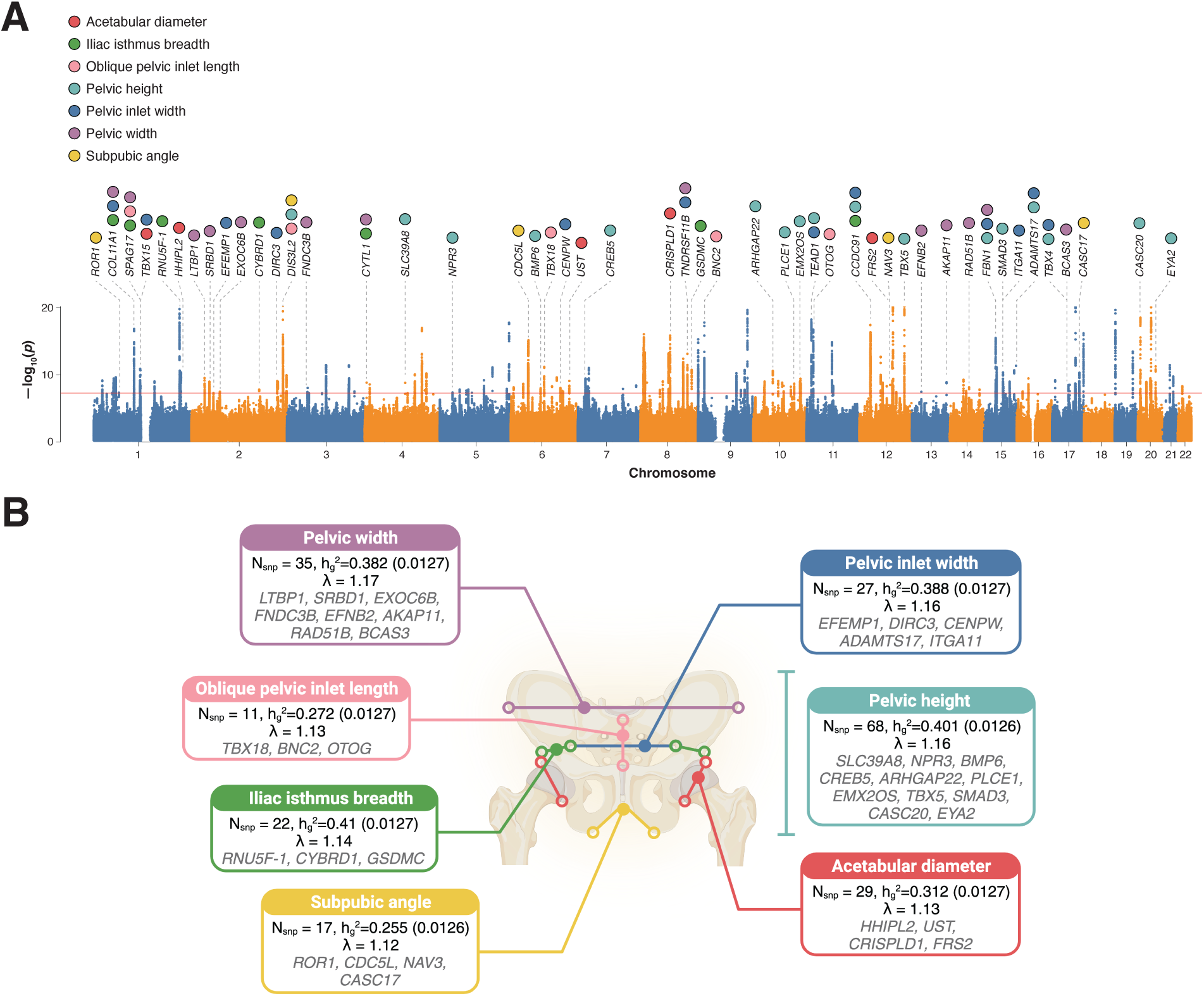
Genome-wide association results. (**A**) Manhattan plot of a GWAS performed across six PPs and subpubic angle; the lowest p for any trait at each SNP is annotated. Loci over the genome-wide significance threshold that are close to only a single gene are annotated. (**B**) Shown are the total number of genome-wide significant loci per trait, heritability (GCTA-REML), λ (from LDSC), and associated genes of loci that are specific to each skeletal trait (again annotating only loci that map to a region with a protein-coding gene within 1000 kb of each clumped region).

### Biological insights from pelvic associations

Out of the 179 independent loci identified across GWASs (**Table S11**), 50 loci overlapped a single protein-coding gene within each clumped region (**Fig. 2B**). Notably, of these 50 genes, 22 (or 44%) resulted in abnormal skeletal phenotypes when disrupted in mice using the Human-Mouse Disease Connection database (*36*). Eight genes (*COL11A1*, *NPR3*, *CDC5L*, *TNFRSF11B, TBX5, FBN1, SMAD3,* and *TBX4*) were associated with rare skeletal diseases in humans (**Table S11**). In some cases, genes associated with specific PPs in our GWAS contribute to human pelvic abnormalities. We found that *TBX15* and *TBX4,* two T-box transcription factors, have been associated with differences in pelvic inlet width and pelvic height in model organisms, and mutations in both the *TBX15* and *TBX4* genes lead to pelvic abnormalities such as hypoplasia of the pelvis and small patella syndrome (*45*, *46*). Thus, our GWAS of PPs identifies genes that were previously associated with skeletal developmental biology and Mendelian skeletal phenotypes and demonstrates the potential for future functional and knockout studies.

### Genetic and phenotypic association of PPs with locomotor phenotypes

We examined how PPs were associated with walking pace, and musculoskeletal disorders such as knee, hip, and back osteoarthritis (OA), which are degenerative conditions that arise from lifetime cumulative effects of gait and motion. First, we used logistic regression to examine phenotypic associations between PPs and these phenotypes (**Fig. 3A**) while controlling for age, sex, weight, height, and other major risk factors for OA (**Methods**: *Polygenic risk score (PRS) association of skeletal phenotypes with musculoskeletal disease*). After correcting for multiple testing at an FDR < 5% across all associations, we found that one standard deviation in two birth canal-related phenotypes was associated with increased self-reported walking pace (oblique pelvic inlet length: *p* = 5.3 × 10^-3^, odds ratio (OR) = 0.96; subpubic angle: *p* = 4.4 × 10^-4^, OR = 0.92) (**Table S15**). As a positive control, we examined another skeletal trait, leg-to-torso length, which we found to be significantly positively associated with walking speed (*p* = 2.97 × 10^-8^, OR = 1.08), in line with previous results and with mechanical modeling (*6*, *7*). These results provide empirical evidence that narrower birth canal proportions in humans are associated with increased walking speed (phenotypic association: between oblique pelvic inlet length and walking pace: *p* = 5.31 × 10^-3^, OR = 0.96, between subpubic angle and walking pace: *p* = 4.89 × 10^-4^, OR = 0.92). However, examining the associations with OA-related phenotypes we found that having larger birth canal-related phenotypes also decreased the risk of back pain/dorsalgia (phenotypic association: between oblique pelvic inlet length and dorsalgia: *p* = 3.45 × 10^-3^, OR = 0.93, between subpubic angle and dorsalgia: *p* = 3.28 × 10^-7^, OR = 0.82, between subpubic angle and back pain: *p* = 5.16 × 10^-7^, OR = 0.87) (**Fig. 3A**, **Table S15**). We also found that individuals with larger birth canal phenotypes were also at increased risk of hip osteoarthritis (phenotypic association: between subpubic angle and hip OA: *p* = 1.18 × 10^-2^, OR = 1.27) but reduced risk of knee osteoarthritis (phenotypic association: between subpubic angle and knee OA: *p* = 9.97 × 10^-4^, OR = 0.83, between subpubic angle and internal derangement of knee: *p* = 9.71 × 10^-5^, OR = 0.81) (**Fig. 3A**, **Table S15**).

**Fig. 3.**
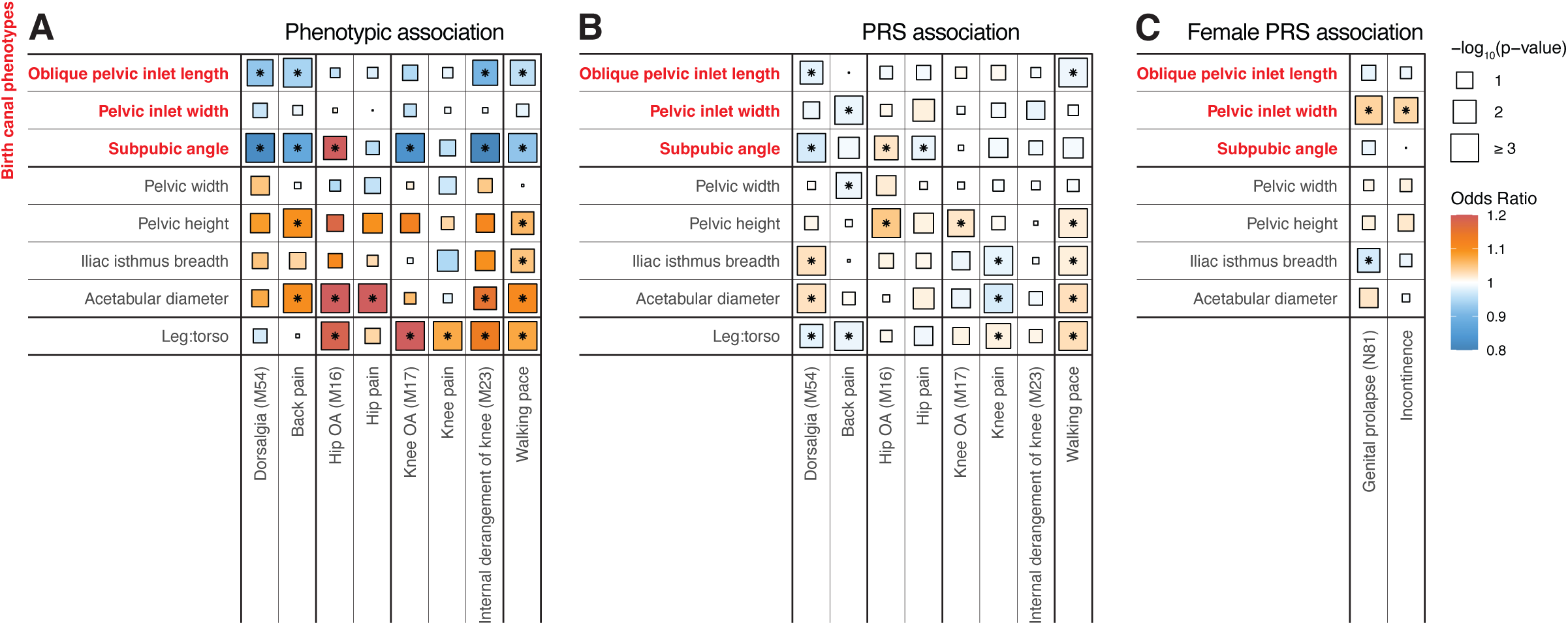
Association between pelvic traits, pain phenotypes, musculoskeletal diseases as well as walking pace. (**A**) Phenotypic associations from logistic regression analyses of musculoskeletal disease traits, self-reported pain and walking pace on PPs. (**B**) PRS associations between musculoskeletal disease traits, walking pace and PPs. (**C**) PRS associations between pelvic floor disorders and PPs. For (A), (B) and (C), associations that are significant after False Discovery Rate (FDR) correction are annotated with an asterisk (*). ORs for the phenotypic associations and PRS are shown in colors, and the p-values are represented by size. The number notations in parentheses are the ICD-10 codes associated with each disease: M54–Dorsalgia, M16–Coxarthrosis (arthrosis of hip), M17–Gonarthrosis (arthrosis of knee), M23–Internal derangement of knee, N81–Female genital prolapse.

To complement these phenotypic associations, we also analyzed 361,140 UKB participants who had not undergone DXA imaging and were of “white British ancestry” for predictive risk based on PRS derived from our GWAS on PPs for the imaged set of individuals (**Fig. 3B**, **Table S16**). We generated PRS with Bayesian regression and continuous shrinkage priors (*47*) using the significantly associated SNPs and ran a logistic regression of the generated risk scores and traits, adjusting for the first 20 principal components of ancestry and imputed sex as well as age, sex, weight and other major risk factors of OA (**Methods**: *Polygenic risk score (PRS) association of skeletal phenotypes with musculoskeletal disease*). Our genetic association analysis mirrored our phenotype association analysis and suggests that individuals with smaller birth canal proportions have on average a faster walking pace, but are at the same time more susceptible to back pain and strain, common consequences of bipedal locomotion due to the distribution of weight on just two limbs (genetic association between leg to torso ratio and walking pace: *p* = 1.00 × 10^-13^, OR = 1.03, between oblique pelvic inlet length and walking pace: *p* = 8.09 × 10^-4^, OR = 0.98, between oblique pelvic inlet length and dorsalgia: *p* = 1.31 × 10^-2^, OR = 0.98, between pelvic inlet width and back pain: *p* = 1.25 × 10^-3^, OR = 0.98, between subpubic angle and dorsalgia: *p* = 1.02 × 10^-4^, OR = 0.97) (**Fig. 3B**, **Table S15**).

### Genetic and phenotypic association of PPs with pelvic floor function

Next, we combined all incontinence-related phenotypes from the ICD10 record, including stress incontinence (N39.3), other specified urinary incontinence (N39.4), fecal incontinence (R15), and unspecified urinary incontinence (R32), into a single binary phenotype. We conducted a GWAS restricted to female individuals who were imaged and computed a PRS for approximately 200,000 females of “white British ancestry” who were independent from the GWAS set. 18,020 individuals out of the 200,000 individuals had one of these incontinence phenotypes. We then regressed binary incidence of genital prolapse and incontinence against PRS for all female pelvic traits, controlling for the number of live births and age (**Methods**: *Polygenic risk score (PRS) association of skeletal phenotypes with musculoskeletal disease*). Out of the various PPs the only significantly positive association we observed was with the width of the birth canal (between pelvic inlet with and genital prolapse: *p* = 4.3 × 10^-4^, OR = 1.04, between pelvic inlet with and incontinence: *p* = 4.2 × 10^-3^, OR = 1.03) (**Fig. 3C**, **Table S17**). These results offer support for the multifactorial pelvic hypothesis, suggesting that a narrower birth canal improves pelvic floor function. Pelvic floor function is critical in assisting bladder and bowel control and evacuation as well as in supporting the fetus during pregnancy - a function thought to be more critical in upright humans than in quadrupeds (*7*).

### Genetic association of PPs with childbirth-related outcomes

Finally, we examined outcomes associated with obstructed labor, which is thought to be more common in humans than any other modern primate species (*9*). Obstructed labor affects around sixteen percent of deliveries today and has been a major cause of maternal and fetal death throughout human history, which suggests it might play a major role in human evolution through natural selection (*48*, *49*). First, we focused on cesarean sections (C-sections) reported in the UK Biobank. To avoid confounding effects due to elective C-sections, we focused on emergency C-sections which are routinely performed in cases of obstruction. We conducted PRS association analysis and found that narrower birth canals were associated with increased risk of emergency C-sections (*p* = 0.0108, OR = 0.92). As childbirth-related outcomes were available only for a small portion of individuals in the UK Biobank (<10% of all individuals) we also examined outcomes in the FinnGen’s data set for delivery-related traits. We identified a significant genetic correlation between birth canal traits and labor obstructions due to maternal pelvic abnormalities, of which a major component is dilation width (**Methods**: *Genetic correlation of skeletal proportions with pregnancy phenotypes*, **Fig. 4B**, **Table S18**). However, we saw no association between obstructed labor due to malpresentation of the fetus and pelvic traits (**Fig. 4B**, **Table S18**). As the position of the fetus can vary independently of the skeletal structure of the pelvis, this childbirth outcome serves as a negative control for this analysis. Combining both types of analysis, our results suggest strong associations between the size of the birth canal and the chance of obstruction during labor.

**Fig. 4.**
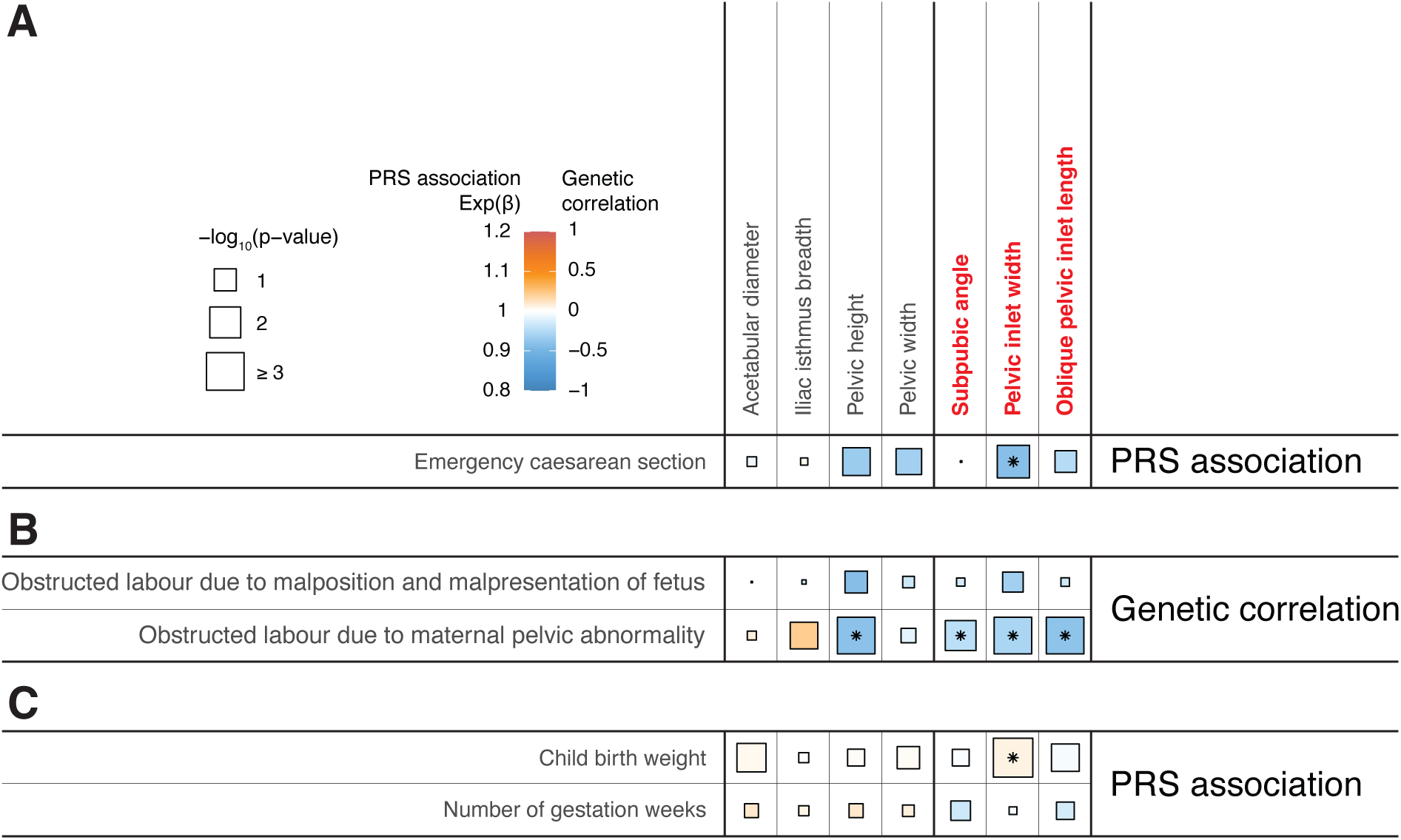
Association between pelvic traits and childbirth-related outcomes. (**A**) PRS associations between pelvic traits and emergency cesarean section. (**B**) Genetic correlations between pelvic traits and obstructed labor, including obstructed labor due to malposition and malpresentation of fetus and obstructed labor due to maternal pelvic abnormality. (**C**) PRS associations between pelvic traits and evolutionary escape variables, including child birth weight and gestation duration. For (A), (B), and (C), associations that are significant after FDR correction are marked with an asterisk. Odds Ratios (ORs) for the PRS associations and genetic correlations are presented in various colors, and the p-values are indicated by size.

### Evolutionary escape

Finally, we investigated associations that might help explain how the obstetrical dilemma may have been alleviated in recent human evolution. First, we examined whether gestation length in humans is shorter than other primates of comparable body size, following from Washburn’s proposed that the relatively large-brained human infant must be delivered before its head reaches a volume that cannot pass through the pelvic canal (*13*, *50*, *51*). However, we found no association between gestational duration and any PP, including those associated with the birth canal (**Fig. 4C**, **Table S17**). This result is in line with more recent data on a fairly large dataset of great apes suggesting that human children are not born significantly earlier than those of the other apes (*14*, *29*–*31*). However, we did see a significant correlation between the proportional width of the birth canal and neonatal birth weight - a proxy for neonatal head size (correlation coefficient ∼ 0.7 (*52*)) (**Fig. 4C**, **Table S17**). This suggests that natural selection might have led to genetic correlation between pelvic and head proportions, potentially reducing labor obstruction (*41*, *53*).

## Discussion

In this study, we used deep learning to understand the genetic basis of skeletal elements that make up human PPs using DXA imaging data in a large population-based biobank. We found sex-specific differences in genetic architecture as well as differences in pelvic symmetry that were associated with handedness. We identified 179 independent genetic loci associated with PPs. We then examined different facets of the obstetrical dilemma, namely the relationship between PPs and locomotor, pelvic floor and childbirth-related outcomes. Lastly, we analyzed possible ways in which evolution and natural selection might have alleviated the dilemma by looking at genetic correlations between gestational period and child birth weight and pelvic proportions.

In previous work on the obstetrical dilemma, studies have examined locomotor outcomes that are associated with efficiency and energy use rather than speed. Here we did not have access to energetics, but we did have access to outcomes associated with walking speed and OA which relate to gait efficiency accumulated over a lifetime. While self-reported walking pace may not seem an ideal measure of walking speed, several lines of evidence suggest that it is a reliable measure of actual walking pace. First, self-reported walking pace is highly heritable (*54*). Second, it is associated with muscle strength and declines with BMI and age in line with expectations (*55*, *56*). It is also associated with several disorders that are known to hinder locomotion, including hip osteoarthritis, the leading cause of adult disability in the United States (*57*–*59*). Finally, self-reported walking pace has been directly correlated with measured walking pace in a reasonable sample size study and within the Biobank (*60*, *61*). It has also been correlated with mean accelerometer assessed activity (*62*).

Our results on locomotion were heavily mixed, with larger birth canal phenotypes related to lower walking speed, reduced risk of back pain and knee OA, but increased risk of hip OA. However, our results provide significant evidence for other facets of the dilemma associated with pelvic floor function and childbirth. Specifically, we show that larger birth canal phenotypes are associated with increased risk of pelvic floor disorders, but at the same time reduced risk of obstruction during labor - two phenotypes that have direct impacts on human evolution due to intense natural selection acting on them. We also investigated several leading hypotheses about how the dilemma could have been alleviated over evolutionary time. Our data does not provide support for the idea that gestational duration has decreased to accommodate birthing large-brained infants - we observed no correlation between any PPs and gestational duration. However, our results indicate that there is a genetic correlation between PPs only related to birth canal width and child head size (which we obtain using birth weight as a proxy phenotype (*52*)). Across all the skeletal traits we examined, the significantly reduced genetic correlation observed between males and females exclusively for birth canal phenotypes also suggests that sexual dimorphism in these traits may have arisen through natural selection in response to different functional constraints.

Beyond increasing the sample size by multiple orders of magnitude relative to previous studies that have examined this hypothesis, and presenting high quality measurement of the human pelvic form, our work is also one of the few studies to integrate data from locomotor, childbirth and pelvic floor outcomes all on the same participants. However, a limitation of our study is that we only had individuals aged between 40-80 years old. It has been suggested that age is a source of variation in PPs, and that changes in functional constraints throughout parts of the reproductive lifespan is another means by which the dilemma could be alleviated (*63*). However, we did not have access to data from individuals from earlier ages to examine this hypothesis.

Taken together, our work combines, imaging, genetic, health record and survey data on biobank scale data to re-examine a 60 year old theory of human evolution that is standardly taught in textbooks. Our results provide major empirical support for several classical theories obstetrical dilemma related to locomotion and childbirth, but perhaps for the first time provides evidence for the role of associated factors such as pelvic floor health.

## Materials and Methods

### UKB participants and dataset

All analyses were conducted with data from the UKB unless otherwise stated. The UKB is a richly phenotyped, prospective, population-based cohort that recruited 500,000 individuals aged 40–69 in the UK via mailer from 2006 to 2010 (*64*). In total, we analyzed 487,283 participants with genetic data who had not withdrawn consent as of May 4, 2022, out of which 42,284 had available DXA imaging data. Access was provided under application number 65439. The baseline participants metadata including age and sex and other variables related to our study are in **Table S2**.

### Dual-energy X-ray absorptiometry (DXA) imaging

The UKB has released DXA imaging data for a total of 50,000 participants as part of bulk data field ID (FID) 20158. The DXA images were collected using an iDXA instrument (GE-Lunar, Madison, WI). A series of 8 images were taken for each patient: two whole body images - one of the skeleton and one of the adipose tissue, the lumbar spine, the lateral spine from L4 to T4, each knee, and each hip. Dual-energy X-ray absorptiometry (DXA) images were downloaded from the UKB bulk data FID 20158. The bulk download resulted in 42,284 zip files, each corresponding to a specific patient identifier otherwise known as each patient’s EID, and each file contained several DXA images of the patient as described above. All images were exported and stored as DICOM files which were later converted to high-resolution JPEG files for image analysis and quantification.

### Phenotype and clinical data acquisition

Self-reported usual walking pace was obtained from UK Biobank under FID 924, and we combined slow pace and steady average pace to increase the sample size in that category. The binary classification of patient disease phenotypes was obtained from a combination of primary and secondary ICD-10 codes (FID 41270) and the non-cancer self-assessment (FID 20002). Self-assessment codes were translated to three-character ICD-10 codes (Coding 609) and ICD-10 codes were truncated to only the initial three characters. Patients received one if a disease code appeared in either self-assessment visit or their hospital records and zero otherwise. The phenotypes related to pelvic floor disorders are derived from ICD-10 codes, specifically including incontinence (stress incontinence ICD-10 code: N39.3, other specified urinary incontinence ICD-10 code: N39.4, fecal incontinence ICD-10 code: R15, and unspecified urinary incontinence ICD-10 code: R32) and genital prolapse (ICD-10 code: N81). As the incontinence phenotypes are very specific and each category of incontinence has only limited data, we combined all incontinence phenotype categories into a single binary phenotype. **Table S12** and **Table S13** contain all ICD-10 and FID codes we used in our analysis.

### Computing infrastructure

All analysis was carried out on the Corral and Frontera system of the Texas Advanced Computing Cluster. The deep learning analysis was carried out on NVIDIA Quadro RTX 5000 GPUs using the CUDA version 11.1 toolkit.

### Image quality control

Each individual in the UKBiobank had a DXA image folder containing up to 8 different body parts. In order to check the labels of these body parts that were defined using their file name, we built a convolutional neural network (CNN) to sort the images by body part through the use of a multiclass classification model using a previously published protocol (*36*). After sorting and removal of images, we were left with 42,228 full skeleton X-rays (**Table S3**). After we determined the final set of full body X-ray images, we performed additional quality control to remove images that were poorly cropped and had other artifacts. To do this we utilized another deep learning classifier also described in (*36*). Removal of all the cropped images resulted in a total of 39,644 full-body images that we used for analysis (**Table S3**).

### Image standardization

From the pool of remaining full-body X-ray images, we discovered that the images varied in both pixel dimension and background. Broadly, the images fell into two main categories: (a) images that were on a black background with sizes between 600-800 by 270 pixels and (b) images on a white background with sizes between 930-945 by 300-370 pixels. The overall distribution of images by pixel ratio and an example of each type of image are shown in **Table S7** and **Fig. S1**. To process these images and remove the effects of scaling and resolution change during the deep learning process, we chose to pad all the images to be of consistent size. We removed images that had sizes far out of the normal range and processed each of the two categories of images separately. The black background images were padded equally on all sides of the image to a final resolution size of 816 × 288 pixels while the white background images were padded in the same fashion to a resolution size of 960 × 384 pixels. We carried this out by converting each individual DICOM file obtained from the UKB into numpy arrays and added additional rows and columns of black or white pixels as appropriate using standard functions from numpy (*65*), scipy (*66*), and skimage (*67*). These final resolution sizes were chosen based on image size requirements for our deep learning model for landmarking and image quantification. Padding and removing individuals with sizes that did not fit into the two major categories resulted in a final total of 39,469 images - 21,981 images of 816 × 288 and 17,488 images of 960 × 384. In our deep learning model for landmarking, we trained using images across both pixel ratios. Despite variations in size and background, the images shared many features, being skeleton X-ray images. Using both pixel ratios enriched the training set, enhancing prediction accuracy.

**Fig. S1.**
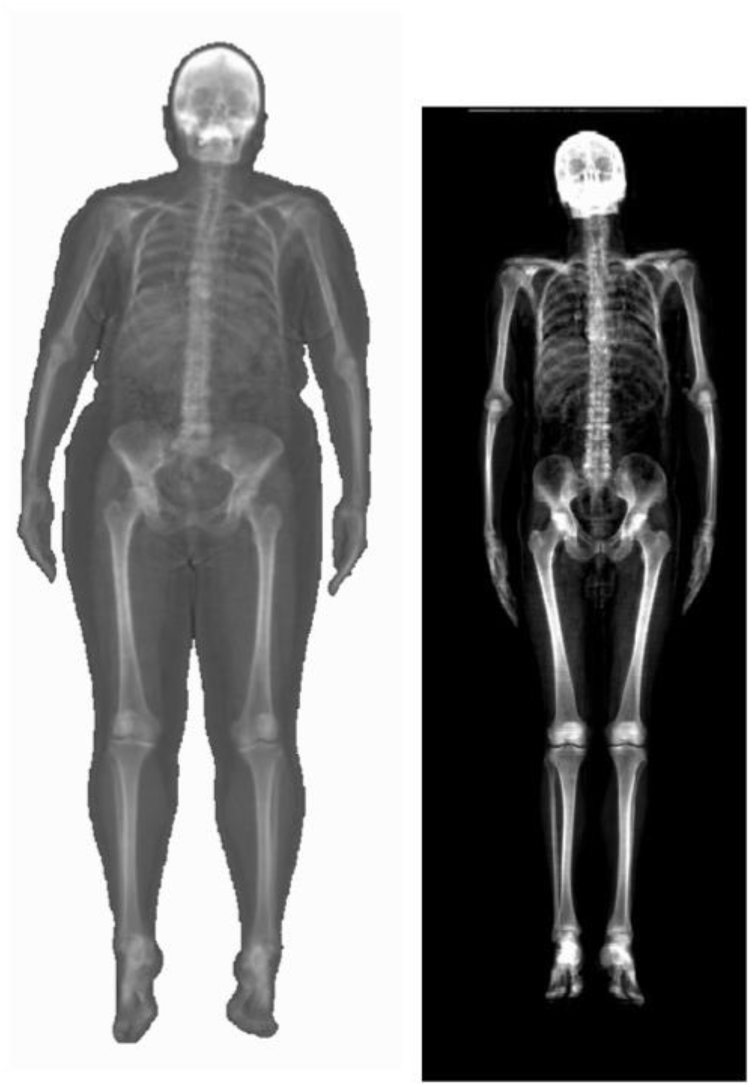
Types of DXA images acquired from the UKB. (Left) Image of patient imaged on white background. (Right) Image of patient imaged on black background. Relative sizes of images are true to scale.

### Manual annotation of human pelvic landmarks

To train our deep learning model, we manually annotated a total of 293 images (with 146 images padded to 960 × 384 pixels on a white background, and 147 images padded to 816 × 288 pixels on a black background). Of these, 239 images were randomly allocated for training and the rest are used for validation. Out of the 293 total images, 10 images were duplicated in each of the image sizes to measure the replicability of our process. We used a single human annotator for all training data and provided an initial dataset of 313 (293+2 × 10 duplicate images) without the annotator’s knowledge. We used a standard annotation scheme in computer vision, the Common Objects in Common (COCO) (*68*) scheme which provides a rubric for joint landmark estimation on the human body. The positions in the pelvis we chose to annotate were the: iliac crest posterior left/right, iliac crest anterolateral left/right, iliac body lateral left/right, pelvic inlet left/right, acetabulum posterosuperior left/right, acetabulum anteroinferior left/rightischiopubic ramus inferior left/right, pubic tubercle, and pubic symphysis inferior. An example of the annotation of one image is shown in **Fig. S2** with landmarks placed at each of the locations listed above.

**Fig. S2.**
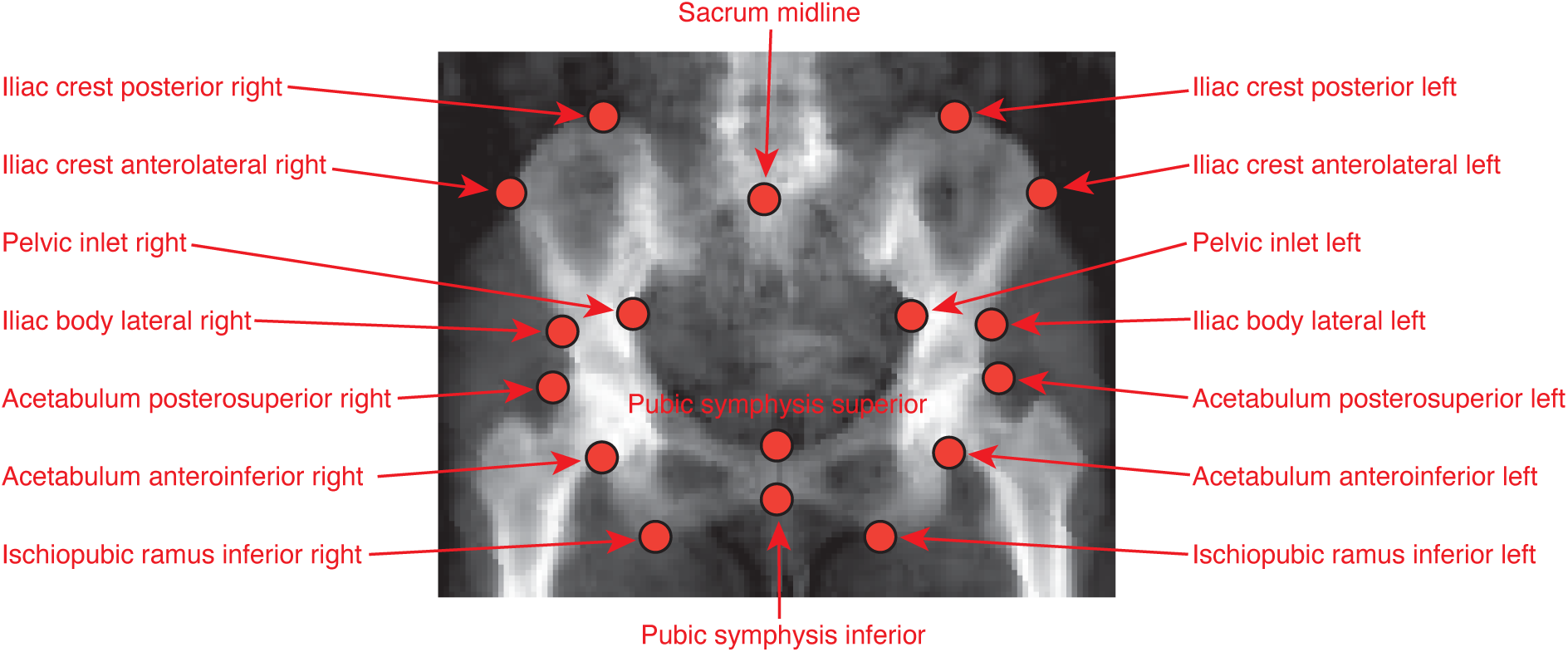
The 17 Pelvic landmarks and their corresponding names.

### Replicability Assessment

We measured the replicability of our annotations by taking the Euclidean distance of pixels between the corresponding key points across 10 randomly selected images that were duplicated amongst both of the 816 × 288 and 960 × 384 image set without knowledge of whether the image was a duplicate. Our replication analysis of 20 duplicate images was under 8 pixels across the different points that were estimated. Across the body parts, the farthest deviation across annotations was seen in the iliac body lateral, but the mean replicability across 20 images was under 2 pixels for all of the pelvic landmarks (**Fig. S3**).

**Fig. S3.**
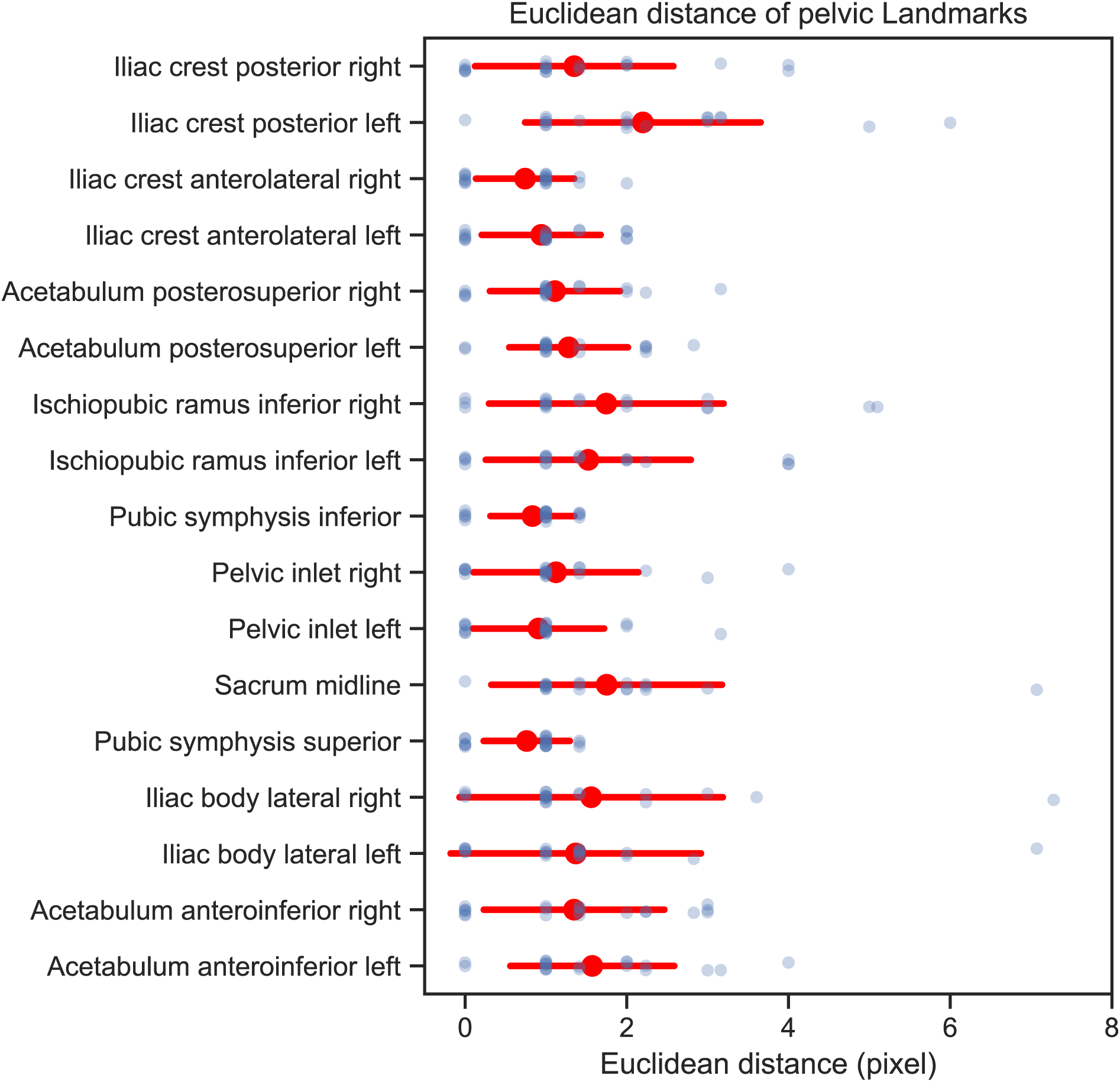
Annotation Error in Human Pelvic Landmarks. The blue points depict the Euclidean distances of specific landmarks between replicate annotations from 10 images of the 816 × 288 set and 10 images of the 960 × 384 set. The red points indicate the mean Euclidean distance for each landmark, while the red error bars represent the standard deviation for these distances.

### A deep learning model to identify pelvic landmarks on DXA scans

To determine the coordinates of all landmarks across 39,469 images, we initially adjusted the DXA images of both sizes, 816 × 288 and 960 × 384, to a uniform dimension of 256 × 256 pixels for the pelvis area by applying central cropping and padding. For the upper and lower body sections, the images were resized to 608 × 608 pixels. Subsequently, 235 images (approximately 80% of the total) were allocated to the training set, with the remaining 58 images set aside for validation. To enhance the training set and to improve the model’s ability to generalize, we applied image transformations including rotation and warping.

To perform landmarking we used an HRNet architecture based network (*69*) preserves high-resolution representations throughout its processing, leading to more accurate estimations (refer to **Fig. S3**). Our previous work demonstrated that using a pre-trained HRNet on large human pose estimation tasks, further refined with fine-tuning, results in more precise predictions (*36*). Consequently, we utilized the pre-trained HRNet architecture (**Fig. S4**), adopting a heatmap size of 64 × 64 for the 256 × 256 pelvis images and 152 × 152 for the 608 × 608 images of the upper and lower body. The batch sizes were set to 16 for pelvis images and 8 for upper and lower body images.

**Fig. S4.**
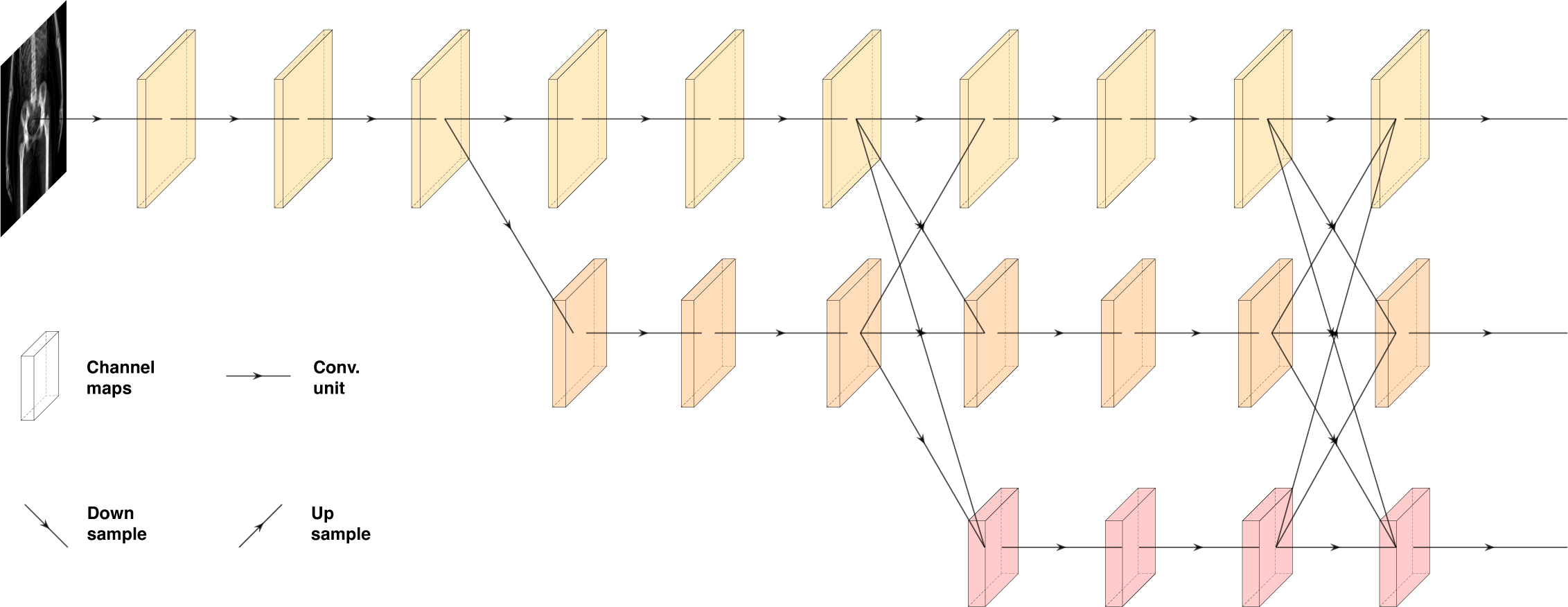
HRNet deep learning architecture. High-Resolution Network (HRNet) architecture maintains parallel high to low-resolution subnetworks.

Initially, our model was trained with 235 manually annotated images. After fine-tuning over 100 epochs using these images and their manually annotated coordinates, we noticed a minimal reduction in loss beyond the 20th epoch (as shown in **Fig. S5**), indicating that 100 epochs were adequate for model convergence. The model’s performance was then assessed using the remaining 20% of annotated images. According to **Fig. S7**, the mean Euclidean distance error between the human annotation and model prediction was below 2 pixels, similar to the error in human annotation (**Fig. S3**). It is notable that certain landmarks, specifically the ischiopubic ramus inferior (left and right) and the iliac crest posterior (left and right), exhibited larger discrepancies between human annotations and model predictions (**Fig. S6**), a variance also observed in repeated manual annotations (**Fig. S3**), highlighting these landmarks as challenging for human annotators as well.

**Fig. S5.**
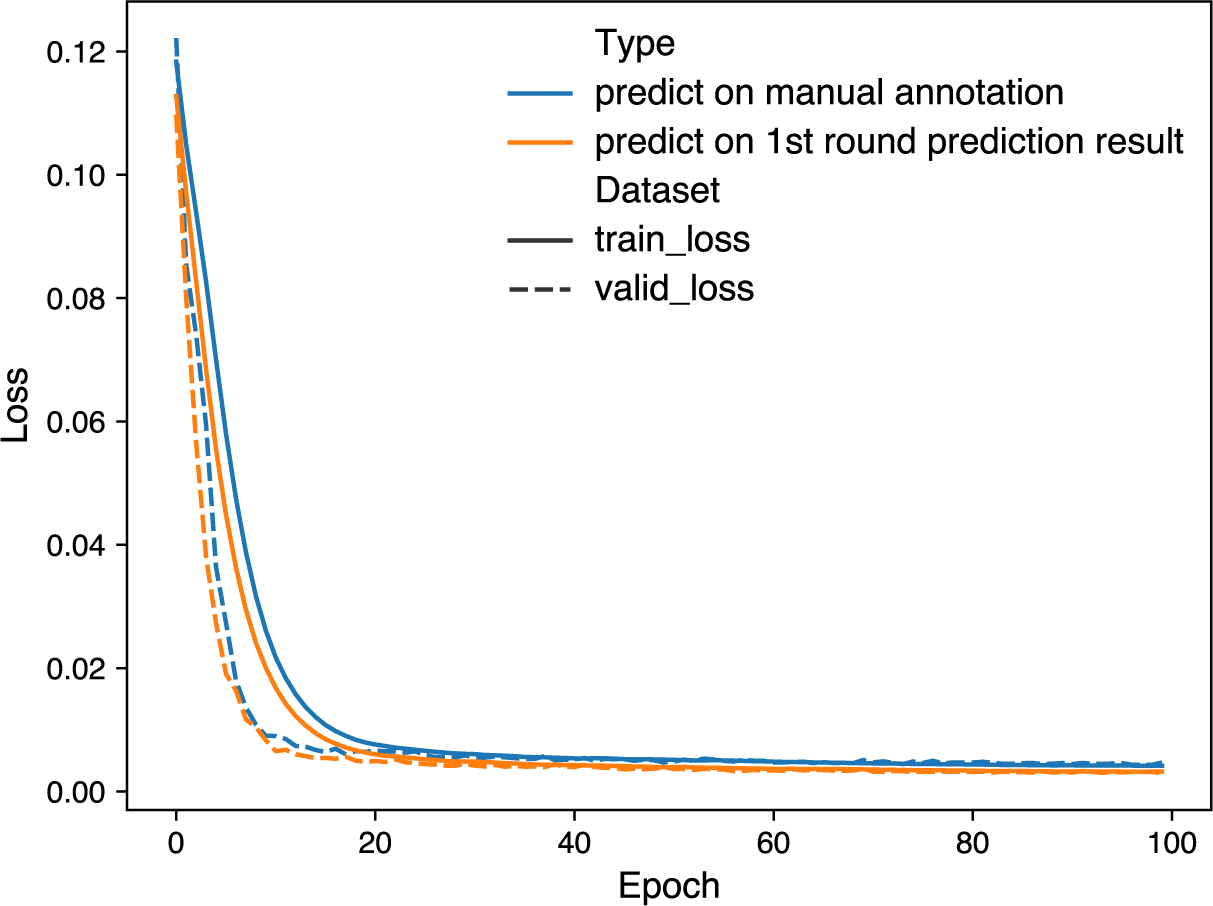
Training logs of HRNet with two different training sets.

**Fig. S6.**
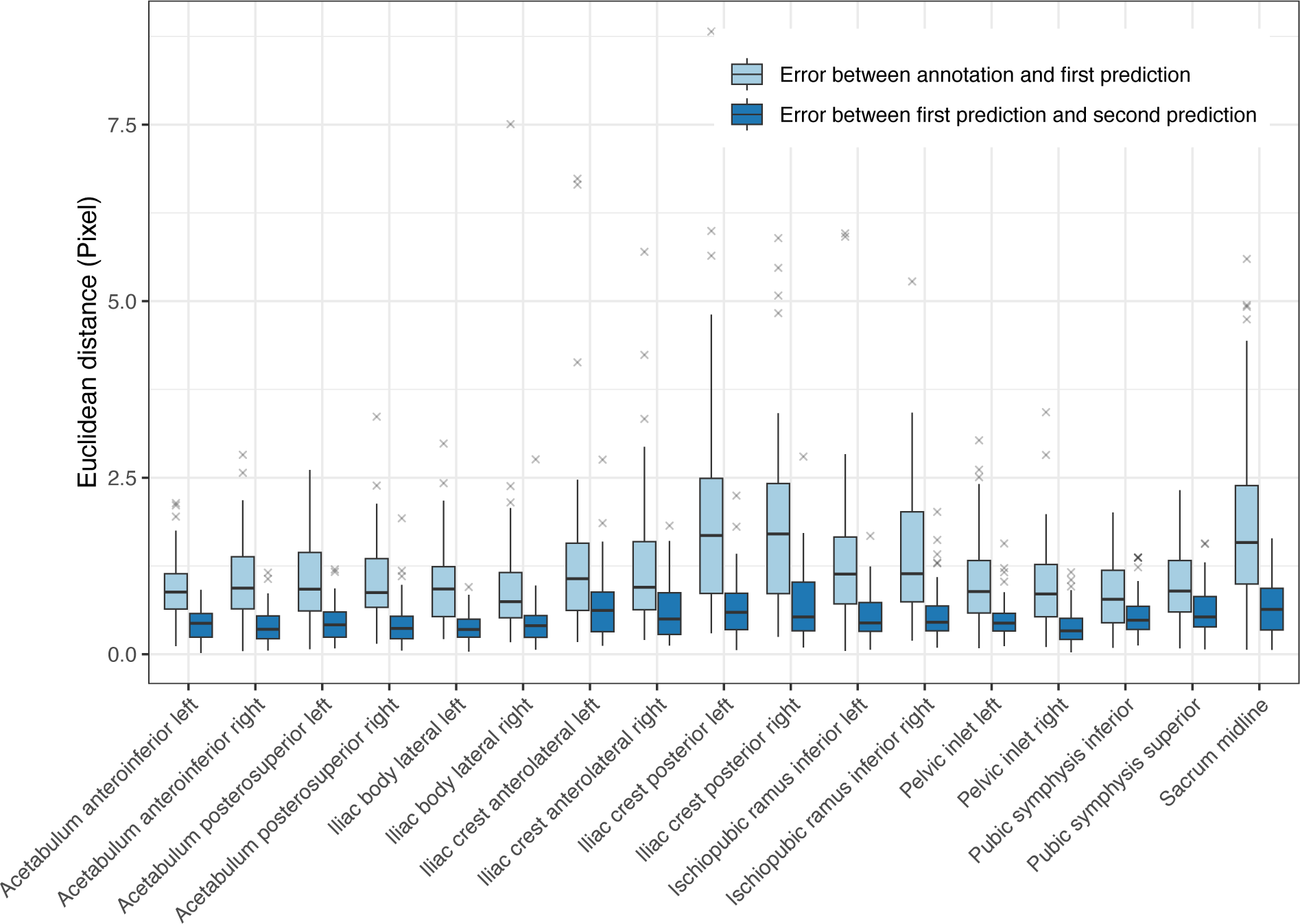
Model prediction error. Light blue box plots indicate the Euclidean distance between manually annotated coordinates and HRNet prediction results based on these coordinates across 18 human pelvis landmarks. Dark blue box plots indicate the Euclidean distance between HRNet prediction results based on previous predicted coordinates across 17 human pelvis landmarks.

**Fig. S7.**
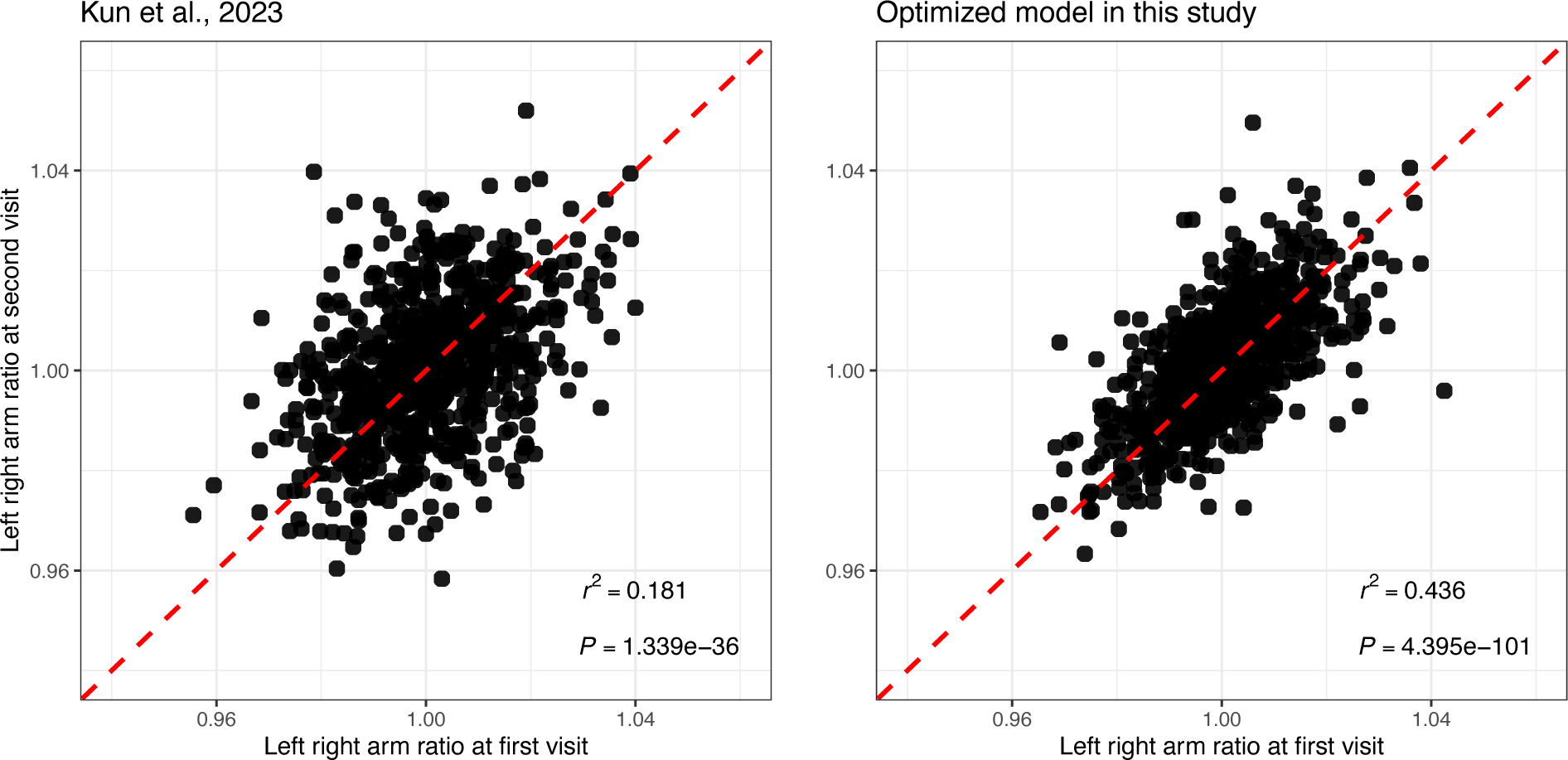
Model Comparison. The left panel displays the scatter plot of the left-to-right arm ratio from two imaging visits using HRNet, sourced from (36). The right panel shows the scatter plot of the same ratio from two imaging visits but using our optimized HRNet model.

We then applied the model to predict landmarks on all remaining images. Initial visual assessments of the model’s predictions on original DXA images suggested greater precision compared to human annotations. To explore this observation, we conducted a validation study, selecting a new set of 293 images from both white and black background sets, ensuring they were not part of the initial training set. These images were split into new training and validation sets, maintaining an 80:20 ratio. Training the model anew with these sets and the same architecture and hyperparameters led to faster convergence and slightly lower loss, suggesting the model’s predicted coordinates might be less noisy than manual annotations (as indicated in **Fig. S7**). Comparisons between human and model-predicted annotations revealed that the Euclidean distance for all 17 landmarks between the first and second model predictions was significantly smaller than the distance between manual annotations and the first model predictions, reducing the average Euclidean distance to less than one pixel (**Fig. 1C**, **Fig. S6, Table S5**).

We further deployed this twice-trained, optimized model on a comprehensive set of 39,469 full-body DXA images from the UK Biobank. Additionally, we compared this model’s performance to that of a previous study’s model (*36*), particularly assessing the correlation between left and right arm length ratios across two imaging visits (**Fig. S7**). Despite using the same annotated coordinates for model training, our current model showed a significantly higher correlation between imaging visits than our previous model. However, we noticed repeated application of this strategy (outlined in **Fig. 1B**) did not yield significant improvements in model accuracy (**Fig. S8**), suggesting that an additional round of training was useful to reduce the variation in manual annotation to a minimum.

**Fig. S8.**
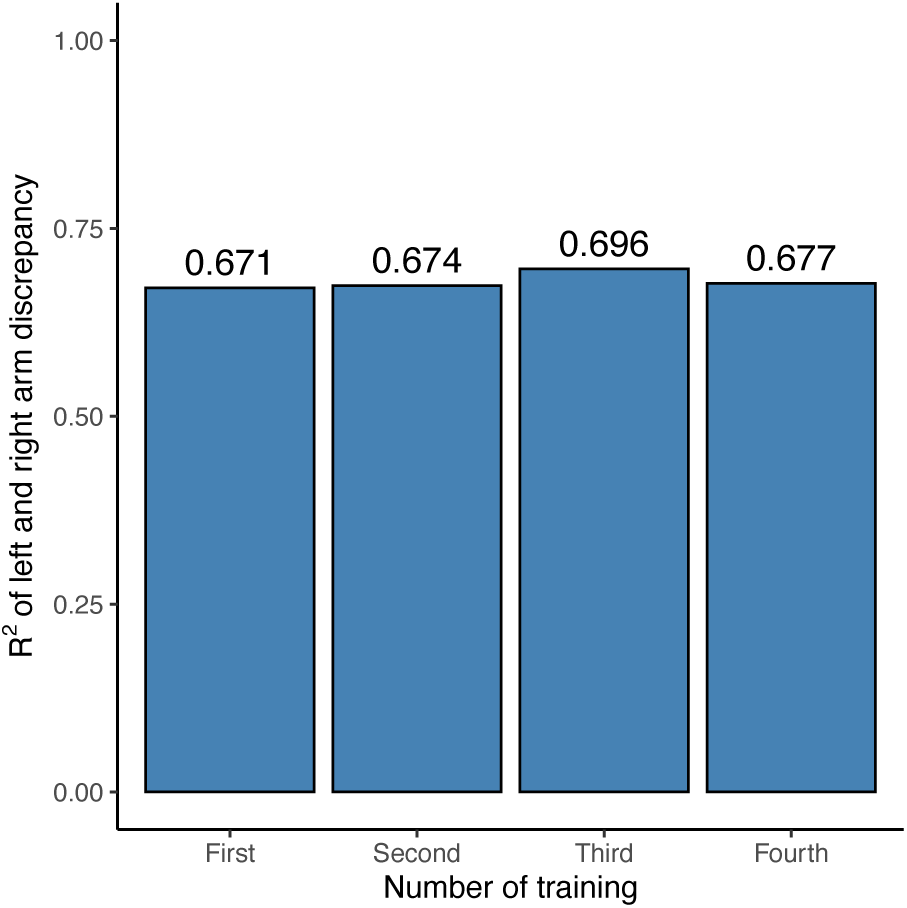
Model prediction accuracy across multiple repeat training.

A major issue in combining our analysis across input pixel ratios was that these pixel ratios represented different resolution scalings, perhaps due to distances that the scanner was held above the patient (**Fig. S9**). That is, in one image a pixel could represent 0.44 cm and in another 0.46 cm. To control for this scaling issue and to standardize the images, we chose to regress height measured directly on our image using the midpoint of the eyes and the midpoint of the two ankle landmarks that could be taken across all image pixel ratios and overall height (FID 50) computed externally from the UKB (**Fig. S10**, **Table S7**). While the height measure we utilized did not include the forehead, it was a relative measure that we used to obtain a scaling factor for each image pixel ratio that we could for normalization. Measurement error of individuals either in our image-based height measure or as reported in the UKB is not expected to affect our conversion from pixels to cms as we are regressing over many individuals. Importantly, we validated this regression and normalization using duplicate individuals taken by different scanners, imaging modes and technicians (**Fig. 1D**, **1E**).

**Fig. S9.**
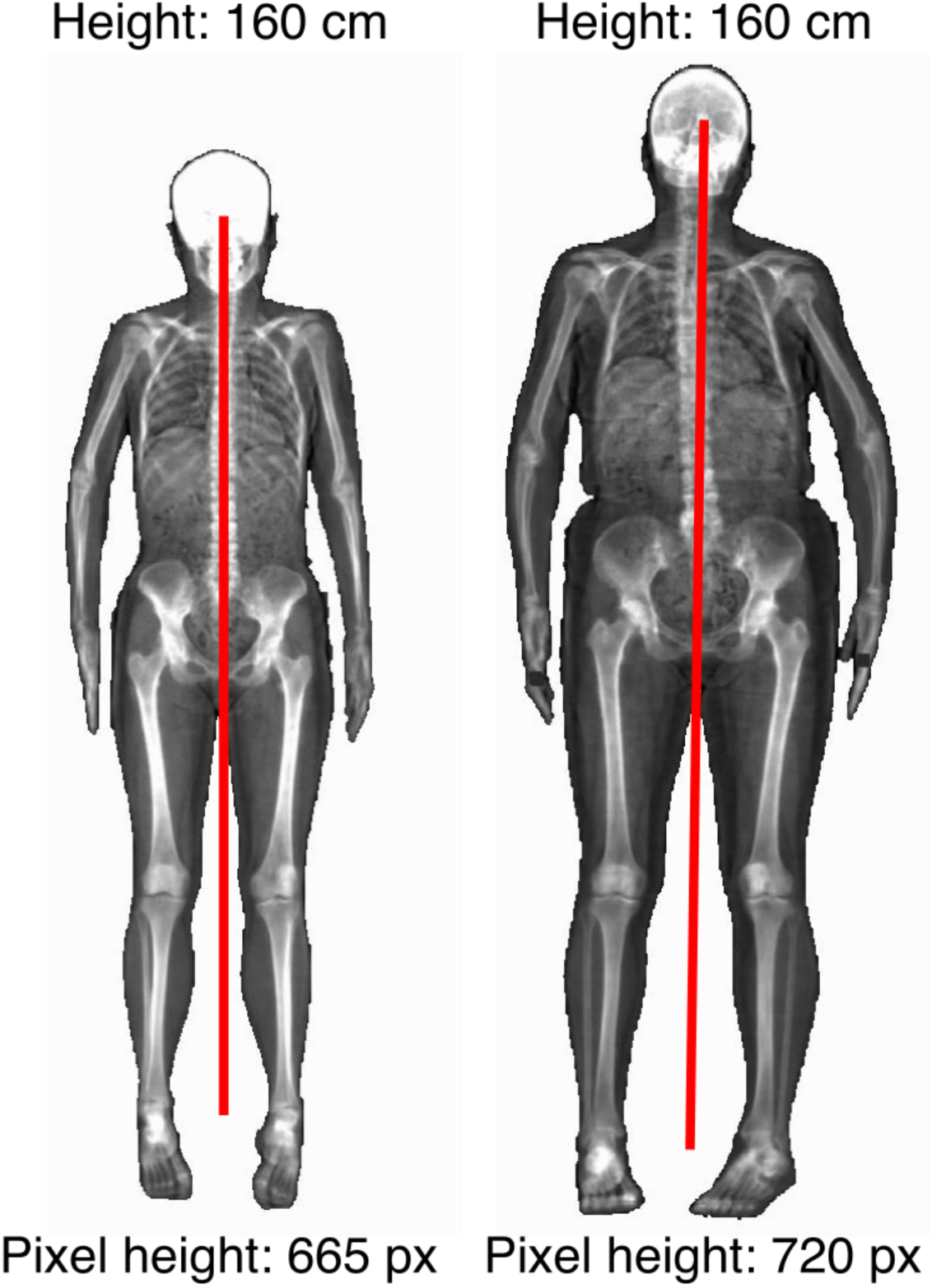
DXA images from the UKB that have undergone different image scaling. Example of two individuals who were measured to be the same height in the FID 50 in the UKB (overall height) but pixel-based measurements of one image were considerably smaller than the other due to image scaling/resolution differences.

**Fig. S10.**
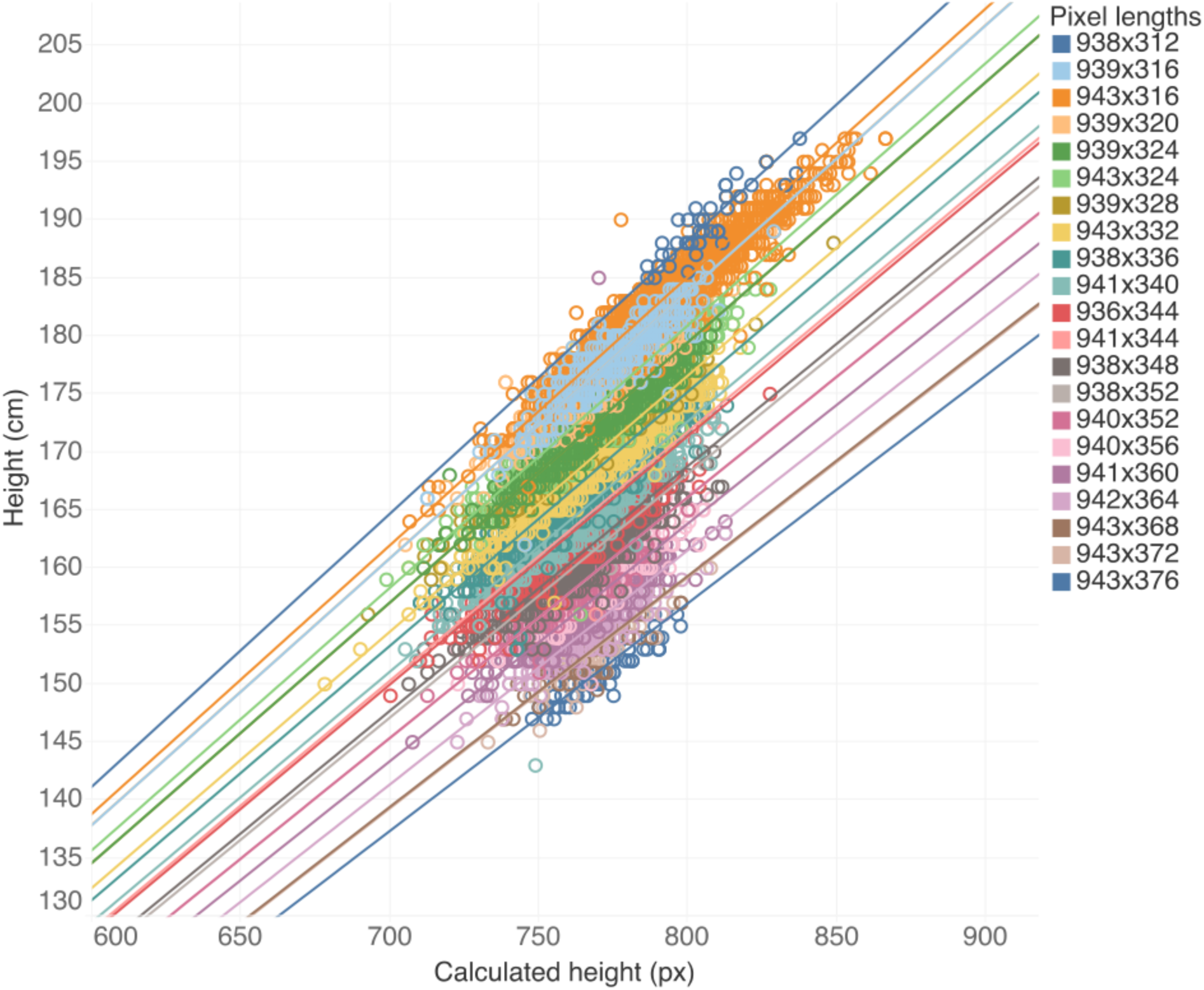
A linear regression of image-measured height against UKB-measured height. For each image pixel-ratio, we regressed height measured in the UKB with height we calculated in pixels from the DXA scan. This provided a conversion from pixels to cm that we used as a normalization factor to correct for differences in resolution.

### Obtaining skeletal element length measures

From each of the 17 landmarks, we generated a total of one angle measure and seven skeletal length measures (**Fig. 1**) in pixels which we converted to centimeters using coefficients from the regressions with height. We averaged iliac isthmus breadth and acetabular diameter across the left and right sides of the pelvis for all analyses. For all measurements, the phenotype values are shown in **Table S8**, and the mean and standard deviations are shown in **Table S9**.

### Participant data quality control

For all of the following analyses, we filtered the participants with correctly labeled full body DXA images (FID 20158 and 12254) to just Caucasian individuals (FID 22006) from the white British population as determined by genetic PCA (FID 21000). We removed individuals whose reported sex (FID 31) did not match genetic sex (FID 22001), had evidence of aneuploidy on the sex chromosomes (FID 222019), were outliers of heterozygosity or genotype missingness rates as determined by UKB quality control of sample processing and preparation of DNA for genotyping (FID 22027), had individual missingness rates of more than 2% (FID 22005), or more than nine third-degree relatives or any of unknown kinship (FID 22021). We also removed individuals whose standing height (FID50) and weight (FID21002) didn’t recorded in their imaging visit. In total 30,370 individuals remained.

### Removal of image outliers

We removed individuals who were more than 4 standard deviations from the mean for any imaging-derived phenotype from the analysis. In total 31,115 individuals remained (**Table S4**). Examination of these outliers by comparing left and right symmetry as well as comparison of other body proportions revealed a heterogeneous set of issues that were associated with the poor prediction by our deep learning model. In some cases, individuals had a limb, or another body part amputated. Some poorly classified images were individuals who had had major hip or knee replacement surgery or had various implants that were causing incorrect model landmark prediction. Another class of outlier images was those that were too poor in quality for any landmarking of any of the points on the image or had abnormal pelvis shape perhaps due to a mendelian disorder (**Fig. S11**).

**Fig. S11.**
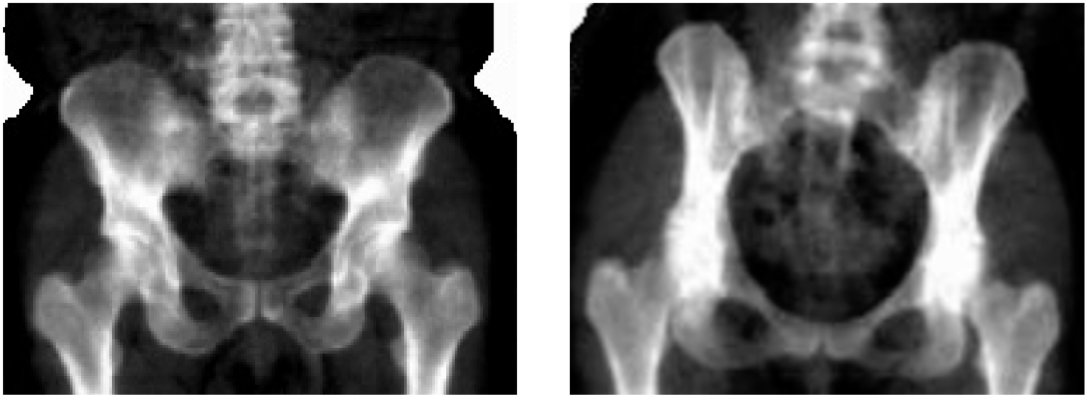
Examples of individuals with normal and abnormal pelvic morphology. Left panel - normal morphology, right panel - pelvis with highly atypical morphology.

### Correlations of pelvic left and right discrepancy with handedness

Upon examining **Fig. 1E**, we observed significant differences between the left and right-side measurements for iliac isthmus breadth and acetabular diameter. Given that the discrepancy between these left and right measurements consistently appeared in two separate imaging visits, this is unlikely to be due to random error/noise in model prediction. To investigate whether this left-right discrepancy is associated with handedness, we calculated the ratio between the left and right-side measurements. We then conducted t-test analyses on the ratio of each phenotype to examine differences in these ratios based on handedness (as indicated by FID1707). These analyses were restricted to white British patients.

### Phenotypic comparison between males and females

For each length phenotype, including pelvic height, pelvic width, iliac isthmus breadth, acetabular diameter, pelvic inlet width, and oblique pelvic inlet length, we regressed out standing height and compared the residuals obtained from the regression analysis between males and females. All six length phenotypes and one angle phenotype showed significant differences between males and females in a t-test (**Fig. S12**).

**Fig. S12.**
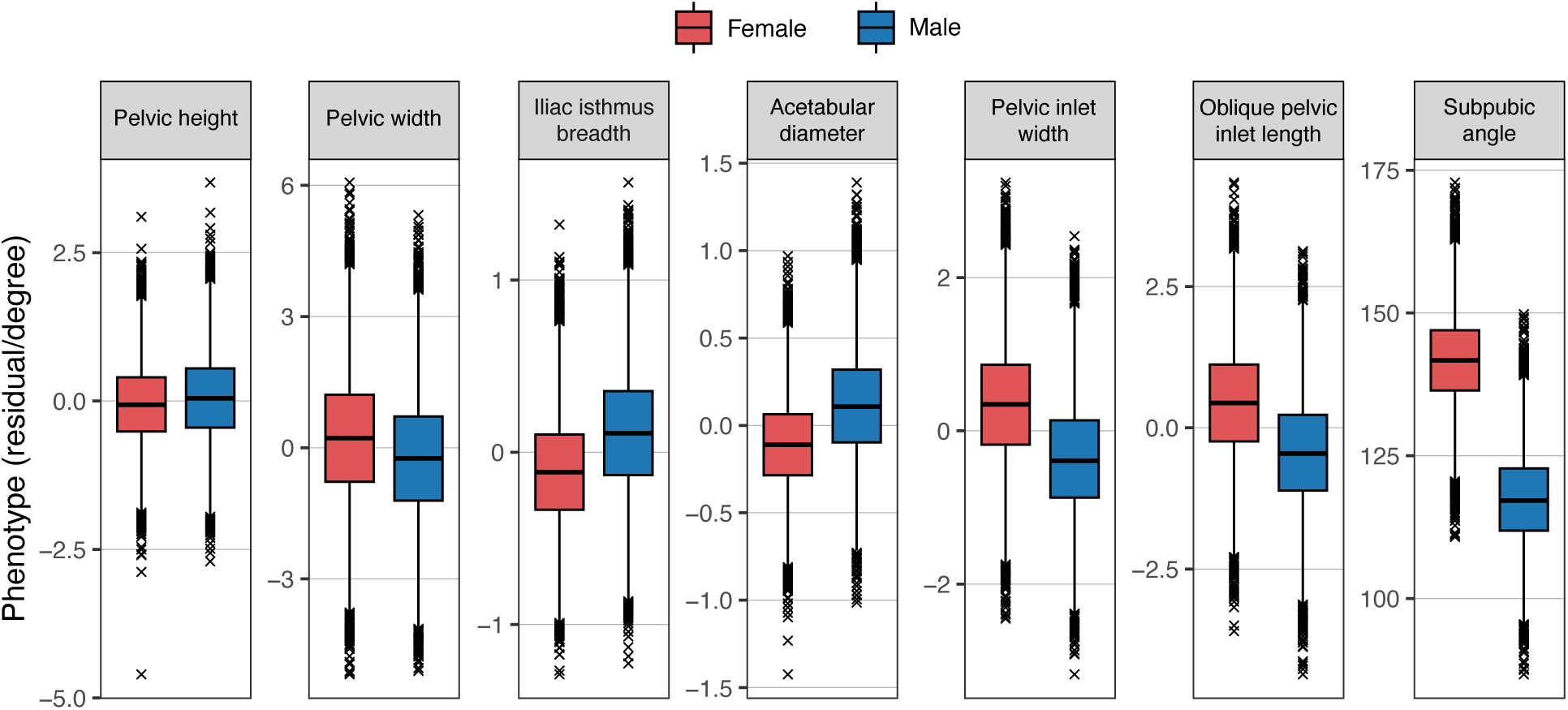
Phenotypic comparison between males and females morphology.

### Genetic data quality control

Imputed genetic data for 487,253 individuals was downloaded from UKB for chromosomes 1 through 22 (FID 22828) then filtered to the quality-controlled subset using PLINK2 (*70*). All duplicate single nucleotide polymorphisms (SNPs) were excluded (--rm-dup ‘exclude-all’) and restricted to only biallelic sites (--snps-only ‘just-acgt’) with a maximum of 2 alleles (--max-alleles 2), a minor allele frequency of 1% (--maf 0.01), and genotype missingness no more than 2% (--maxMissingPerSnp 0.02). In total 8,638,168 SNPs remained in the imputed dataset. Non-imputed genetic data (genotype calls, FID 22418) did not contain duplicate or multiallelic SNPs but were filtered to the quality-controlled subset; 652,408 SNPs remained.

### Adjusting for height correlation in GWAS by adding height as covariate

A major issue in carrying out GWAS for phenotypes such as ours where we would like to control for height is the potential for confounding due to the adjustment. McCaw et al. highlight the pitfalls in GWAS of ratio traits and describe ways to reduce this confounding (*43*). Following their pipeline, we carried out GWAS, adjusting not only for height but also for leave-one-chromosome-out (LOCO) polygenic scores (PGS) of height. Briefly, we first performed GWAS on approximately 370k white British individuals without imaging data using PLINK (*70*). Second, we estimated LOCO-PGS for each individual with imaging data for each chromosome using Bayesian regression with continuous shrinkage priors (*47*), employing associated single nucleotide polymorphisms from HapMap3. In total, each individual received 22 LOCO-PGS one for each chromosome. Third, we randomly split 30k imaged individuals into two groups, with each group containing around 15k individuals, and conducted two GWAS models for each chromosome. These models adjusted for the first 20 principal components, age, sex, height, and, optionally, the corresponding LOCO-PGS as covariates. Finally, we concatenated all chromosomes’ GWAS summary statistics. In examining the effect sizes across all manners of carrying out the analyses, we found that results (*p* < 5 × 10^-4^) adjusted for and not adjusted for LOCO-PGS were very similar in the split-sample study (**Fig. S13**), suggesting that the genetic component of collider bias is minimal. In **Fig. S12** we present an image of the correlation of effect sizes for only a single trait but the results of all traits ranged in correlation between 0.959 to 0.989. To provide additional confirmation of reduced confounding with the adjustment for height, we compared the effect size correlation between snps at *p* < 5 × 10^-4^ for height and a specific trait (without adjusting for LOCO-PGS) and observed that effect sizes estimated for height and the SNP effect size of a specific trait were completely uncorrelated (**Fig. S14**). This further indicates that the results of the GWAS we conduct for particular pelvic proportions are largely independent of height.

**Fig. S13.**
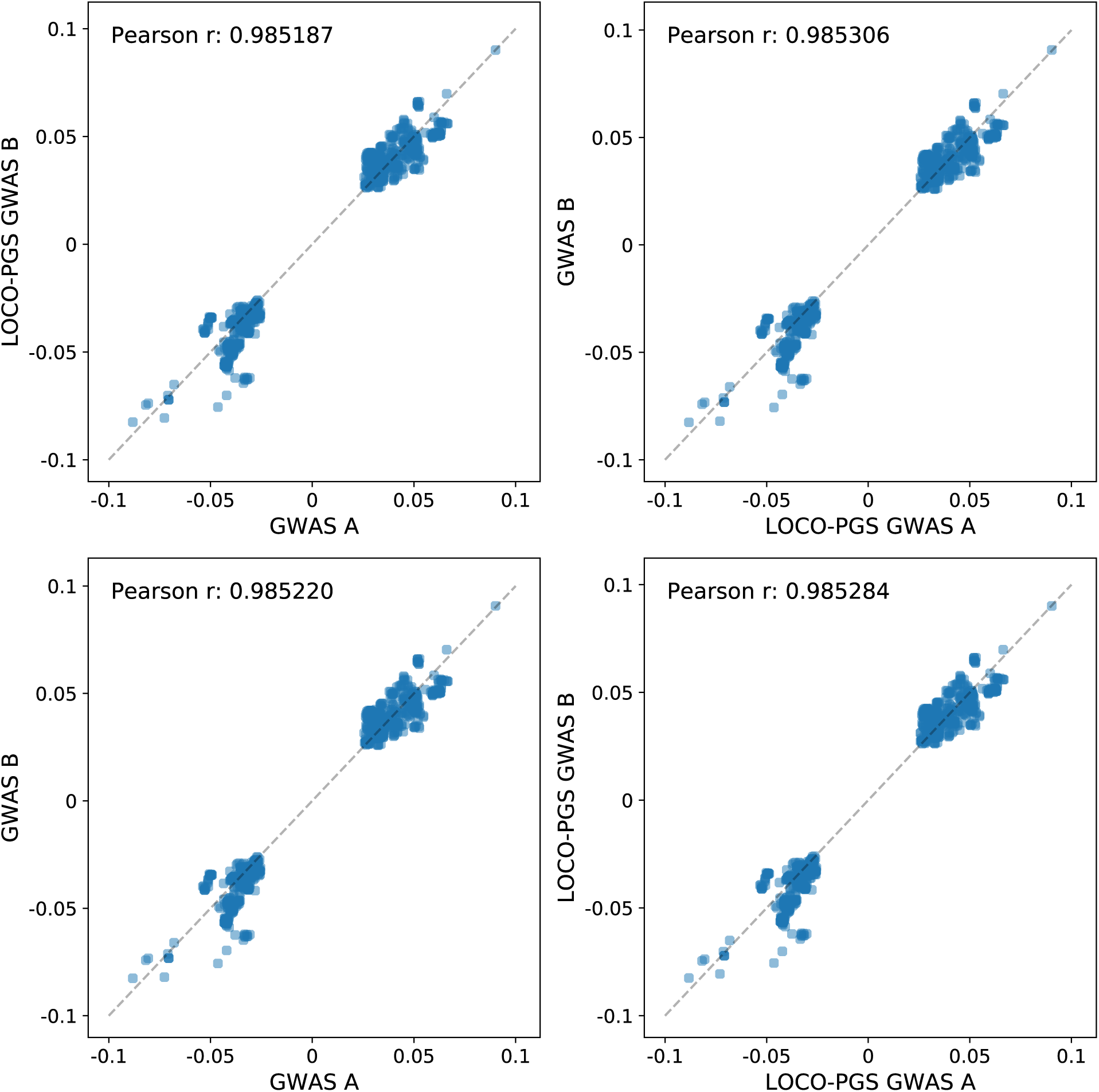
Two GWAS model comparison. To compare with or without adjusting for LOCO-PGS we compared the effect size correlation between two separate samples with the same or different GWAS models with SNP p < 5 × 10^-4^. Here we only show one randomly picked trait birth canal width, but we observed similar signals for all traits.

**Fig. S14.**
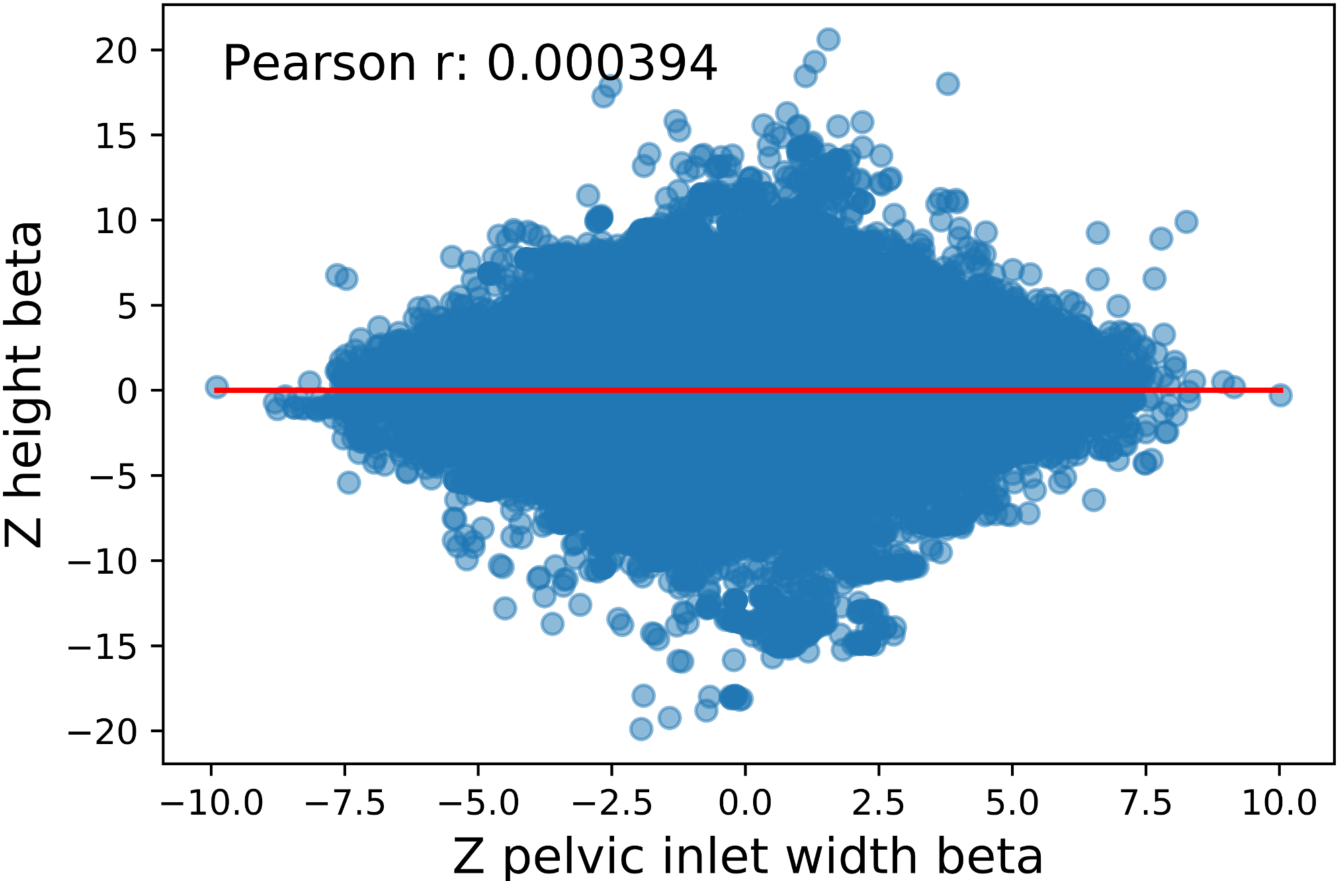
SNP effect size correlation between height and birth canal width.

### GWAS and Heritability analysis

GWAS was performed with BOLT-LMM (*47*). LD Score v1.0.1 was used to compute linkage disequilibrium regression scores per chromosome with a window size of 1 cM (*44*). PLINK2 --indep-pairwise with a window size of 100 kb, a step size of 1, and an r^2^ threshold of 0.6 was used to create a list of 986,812 SNPs used as random effects in BOLT-LMM. Covariates were the first 20 genetic principal components provided by UKB (FID 22009), sex (FID 31), age (FID 21003), age-squared, sex multiplied by age, sex multiplied by age-squared, and standing height (FID 50). In addition, the DXA scanner’s serial number and the software version used to process images were combined into one covariate, resulting in 5 factor levels.

SNPs in each resulting GWAS were clumped in PLINK using --clump with a significance threshold of 5.0 × 10^-8^, a secondary significance threshold of 1.0 × 10^-4^ for clumped SNPs, an r^2^ threshold of 0.1, and a window of 1 Mb. SNPs were assigned to genes with -- clumpverbose --clump-range glist-hg19 downloaded from PLINK gene range lists (*71*). The genomic inflation factor of each phenotype was assessed in R version 4.2.1 as the ratio of the median of the observed chi-squared distribution (an output of BOLT-LMM --verbose) to the expected median of the chi-squared distribution with one degree of freedom.

We created the genetic relationship matrix for our quality-controlled subset but without any related individuals and a minor allele frequency of 0.01, then ran GCTA for each phenotype pair with the first ten genetic principal components provided by UKB (FID 22009).

The heritability of each phenotype was assessed with European HapMap3 SNPs using GCTA (*38*) with the same covariates as GWAS, excluding age-squared and sex by age-squared. We also estimated heritability using LDSC (*44*) and found similar heritabilities (20-50%) (**Fig. S15**, **Table S10**).

**Fig. S15.**
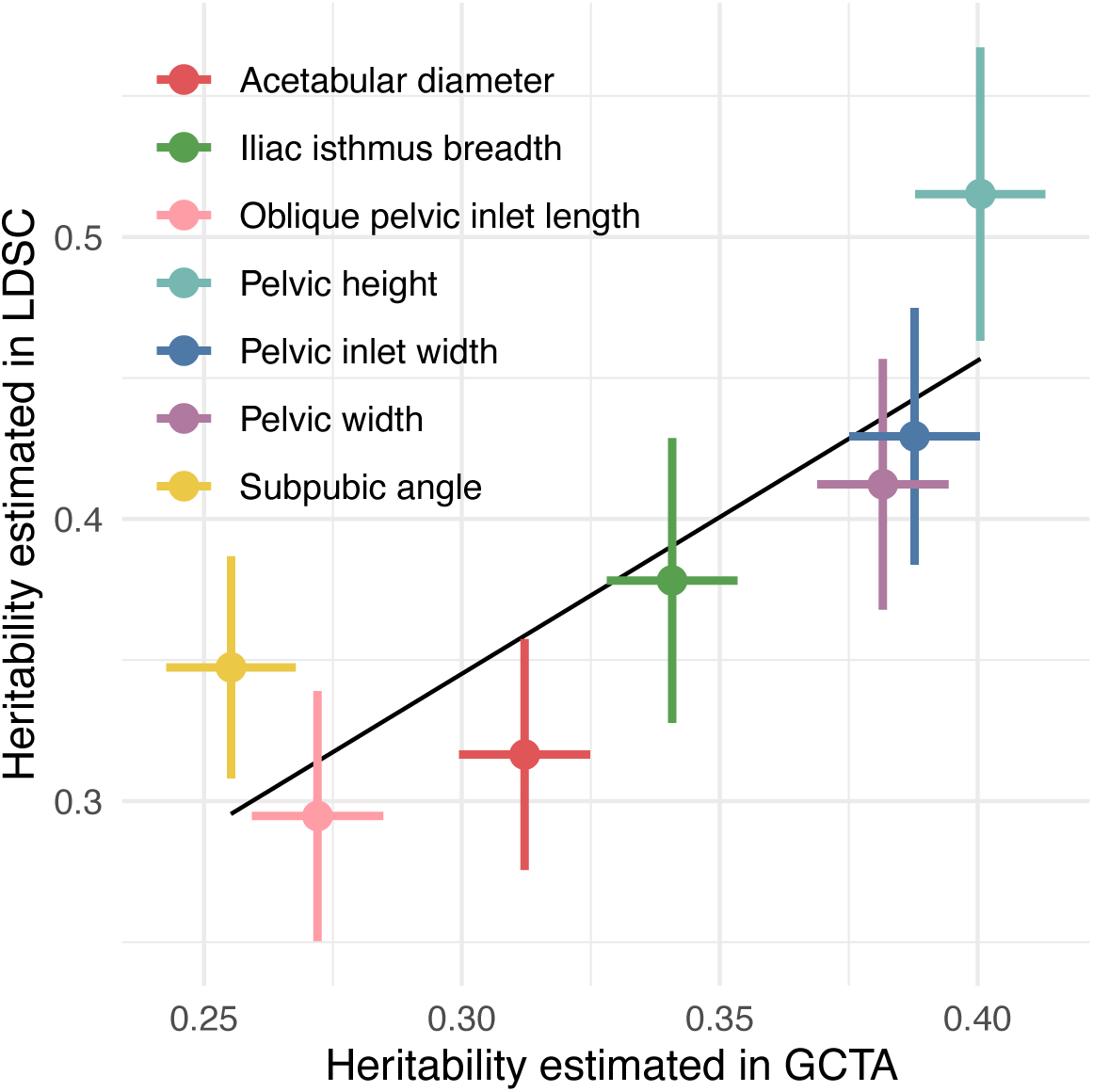
Heritability estimated in GCTA and LDSC.

### Sex-specific analysis

We performed a GWAS independently in males and females using the same process and covariates we used in the combined GWAS analysis in the previous section. Subsequently, we used LDSC to carry out genetic correlation analysis between GWAS conducted in males and females. As depicted in **Fig. 1F**, outer pelvic morphology, such as pelvic height, had genetic correlation consistent with 1. However, phenotypes related to the birth canal, such as pelvic inlet width, oblique pelvic inlet length, and subpubic angle, exhibit differences significantly different from 1. This aligns well with previous studies, underscoring the functional importance in females to accommodate childbirth.

To determine if any sex-specific loci were present in our pelvic phenotypes, we also carried out additional GWAS in PLINK involving a Sex-Genotype interaction for each SP on our original population of 31,115 individuals to determine loci with sex-specific effects. Across all the traits that we examined we did not find evidence for interaction at any locus which would signify sex-specificity. However, we note that this lack of evidence could possibly be due to reduced power for detecting interaction effects at this sample size. We also report the summary statistics for this GWAS with interactions along with the other GWAS that we performed in the Supplementary Data.

### Clumping and identification of genes associated with loci

To obtain a set of independent SNPs associated with each PP phenotype, we first performed clumping analysis for each phenotype using plink and assigned SNPs to genes with -- clump-verbose --clump-range glist-hg19 with an r^2^ window of 0.1 and a 1 Mb threshold of physical distance for clumping. We downloaded gene ranges from plink for hg19 (*72*). Following clumping, we looked at a subset of 7 phenotypes and combined the significant SNPs across the chosen phenotypes resulting in 339 unique SNPs.

### Functional mapping and gene enrichment analysis

We ran FUMA (*73*) without any predefined lead SNPs on a sample size of 31,115 individuals. GENE2FUNC was run with all types of genes selected as background genes using Ensembl v92 with GTEx v8 gene expression data sets and we set window sizes 10 kb for both upstream and downstream (**Fig. S16**, **Table S19**).

**Fig. S16.**
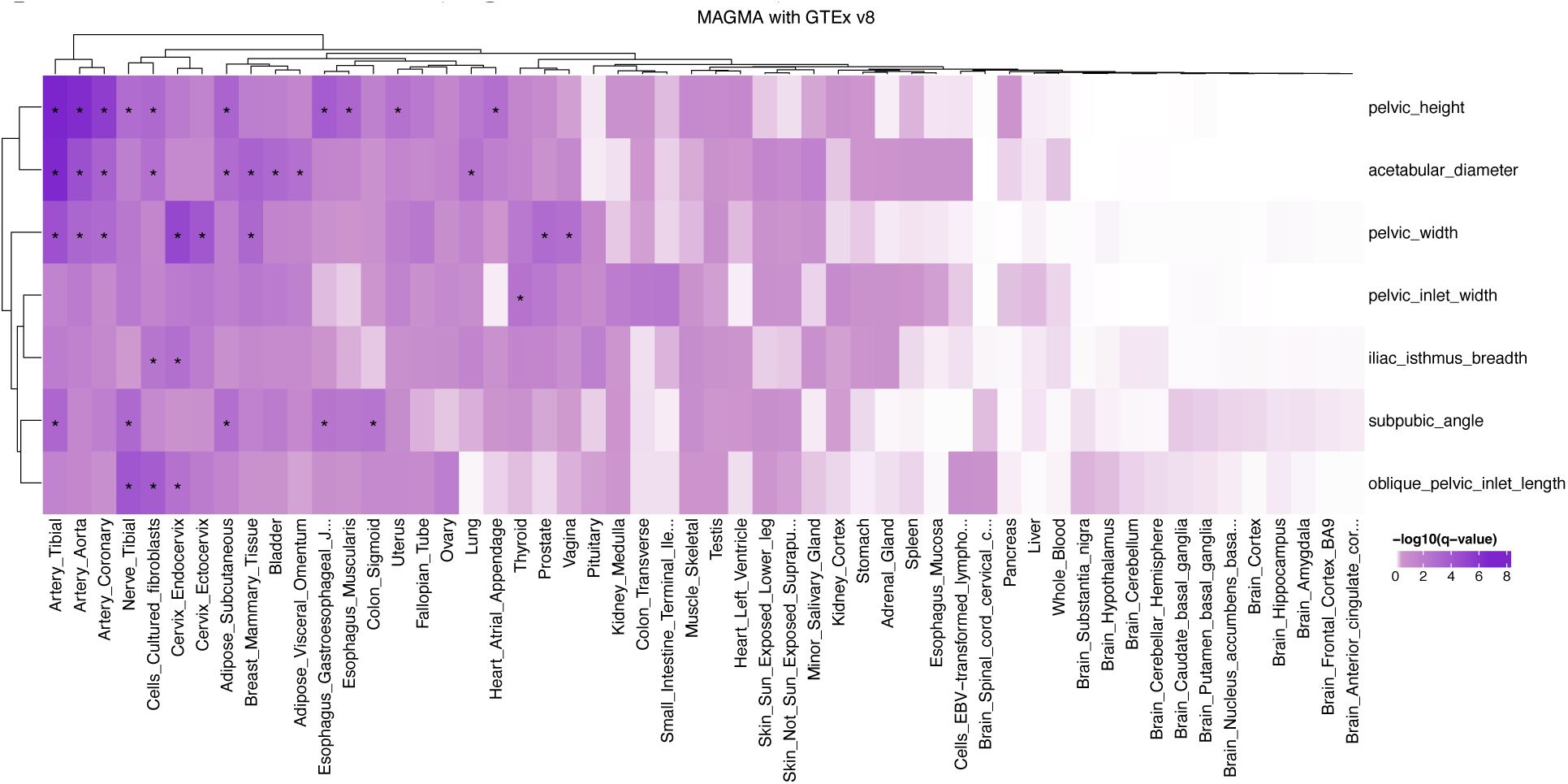
MAGMA gene property analysis with GTEx 8 and image-derived phenotypes GWAS.

### Transcriptome analysis

To explore the genetic underpinnings of pelvis-related phenotypes in relation to specific developmental stages of the human pelvis, we sought enrichment of genes associated with our GWAS results for pelvis-related phenotypes in gene expression data across four distinct developmental stages of the human pelvis during the embryonic period, as detailed in (*33*). Our primary objective was to discern which developmental stage (E53, E54, E57, or E59) might be linked to changes in pelvis shape. We downloaded RNA-Seq data for human embryonic pelvises at different developmental stages from the GEO data repository (GSE165930). Subsequently, we converted gene names to Ensembl gene IDs using the biomaRt package (version 2.52.0) in R. To compute the relative gene expression level for a specific subelement at a particular developmental stage, we subtracted the average expression from other stages for that specific subelement and from other subelements across different stages. Following this, we conducted a MAGMA gene property analysis to assess enrichment between genes expressed during specific developmental stages and our phenotypes. However, our analysis did not reveal any significant enrichment for any developmental stage in our GWAS after adjusting for multiple comparisons using FDR correction for both the number of traits and developmental stages (**Fig. S17**, **Table S20**). In a subsequent approach, we combined data from different developmental stages to investigate potential associations between pelvis-related phenotypes and specific pelvis subelements. We determined the relative expression of specific subelements by subtracting the average expression of other subelements. Another round of MAGMA gene property analysis revealed a significant effect between the Ilium and pelvic inlet width, as well as between the Acetabulum and subpubic angle, after FDR correction (**Fig. S18**, **Table S21**).

**Fig. S17.**
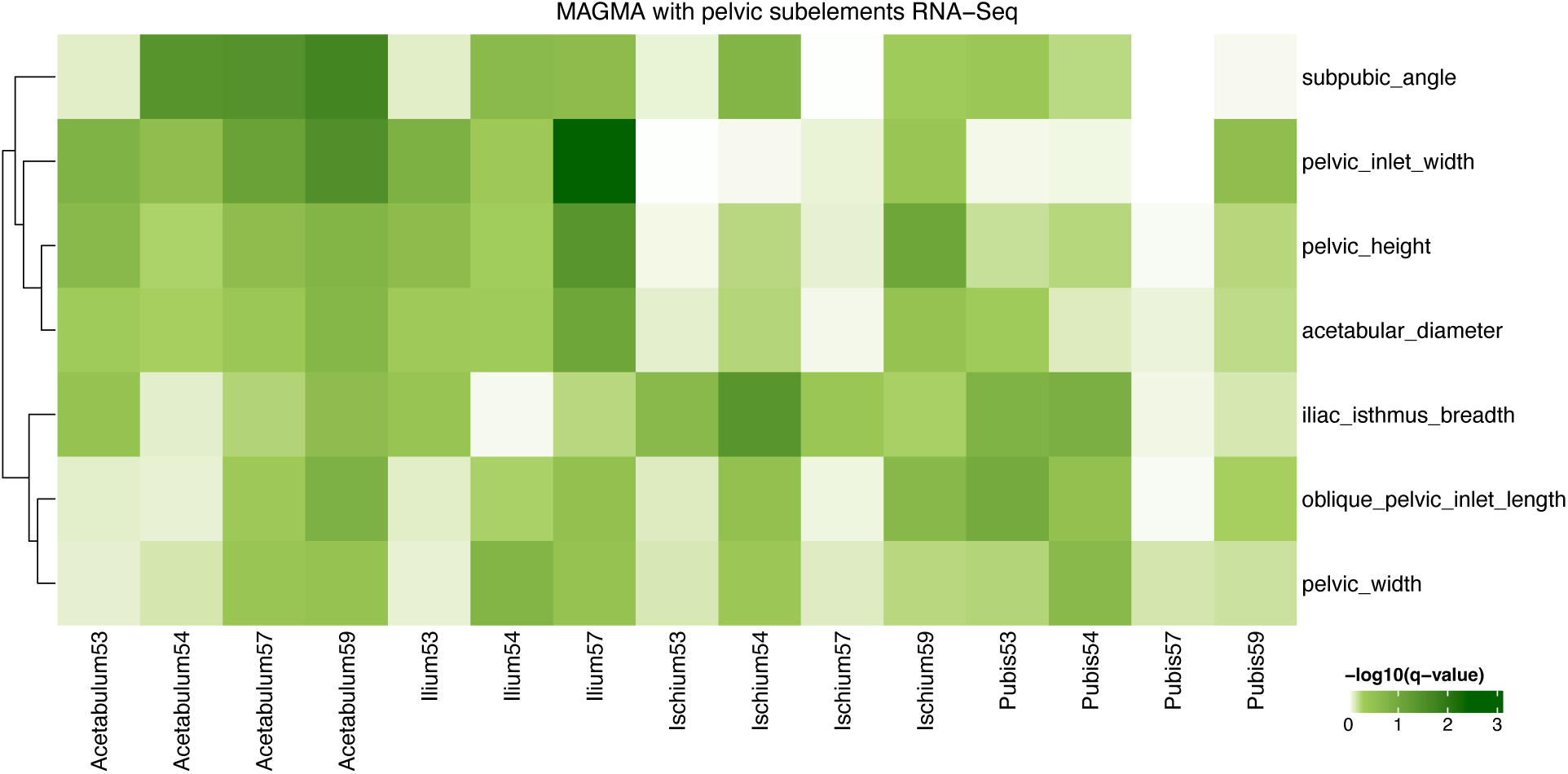
MAGMA gene property analysis with pelvis subelements in different developmental stages ATAC-Seq and image-derived phenotypes GWAS.

**Fig. S18.**
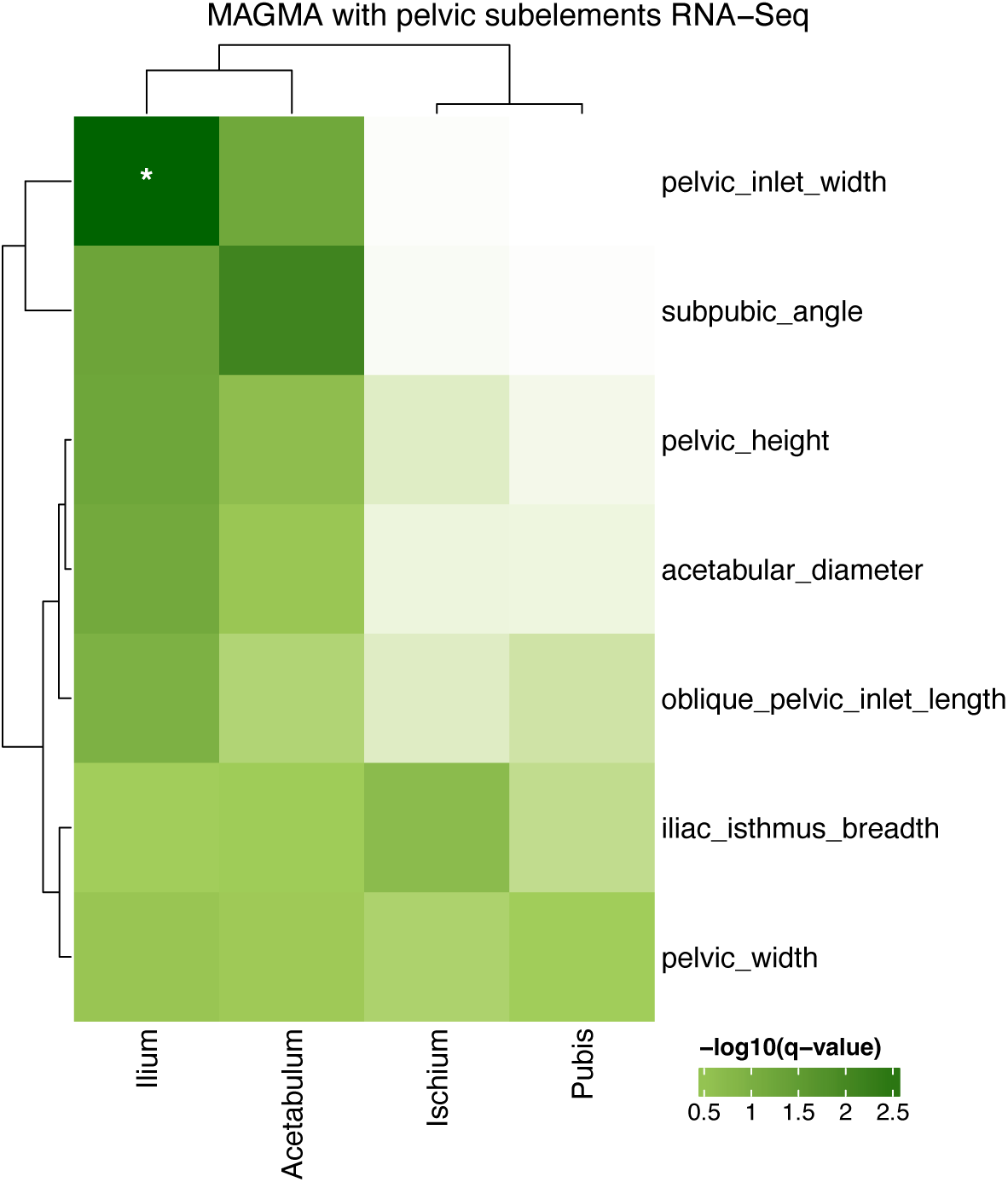
MAGMA gene property analysis with pelvis subelements ATAC-Seq and image-derived phenotypes GWAS.

### Phenotypic association of skeletal phenotypes with musculoskeletal disease

To examine correlations between our pelvis phenotypes with musculoskeletal disease, musculoskeletal or connective tissue diseases related to the hip, knee, and back we obtained data from UKB Chapter XIII (FID 41270) ICD-10 codes as well as self-reported pain phenotypes (FID 6159) for the hip, knee and back. We then regressed the binary outcome of disease or reported pain against pelvis phenotypes controlling for clinically relevant covariates that are known to affect OA (*74*) including age, sex, diet, BMI, and other factors. A full list of variables we controlled for are reported in **Table S14**. After running the regressions, we used Bonferroni correction for significance at the level of the total number of disease/pain traits multiplied by the total number of skeletal phenotypes.

### Polygenic risk score (PRS) association of skeletal phenotypes with musculoskeletal disease

This analysis only utilized the ∼370,000 white British individuals who were not included in our imaging dataset for which GWAS was conducted. We generated PRS for each of the generated traits with Bayesian regression and continuous shrinkage priors (*47*) using the associated single nucleotide polymorphisms from HapMap3. We ran a logistic or linear regression of the PRS on traits across all individuals, adjusting for weight, household income, non-insulin-dependent diabetes mellitus (ICD-10 code: E11), depressive episode (ICD-10 code: F32), recurrent depressive disorder (ICD-10 code: F33), chronic ischaemic heart disease (ICD-10 code: I25), smoking status (FID 20116), and sleep duration (FID 1160). For female PRS regression we also adjusted for the number of live births (FID 2734).

### Genetic correlation of skeletal proportions with pregnancy phenotypes

We utilized cross-trait LD score regression (https://github.com/bulik/ldsc/wiki/Heritability-and-Genetic-Correlation) for estimating genetic correlations between each of our pelvis-related phenotypes and case-control pregnancy phenotypes from the Finngen (https://www.finngen.fi/en/access_results) by using GWAS summary statistics.

## Supplementary Tables

**Table S1 *-*** Previous studies have attempted to test the obstetrical dilemma hypothesis.

This table contains the papers involved in the debate on the obstetrical dilemma.

**Table S2 -** GWAS population summary

This table contains summary data on the population subset used in our GWAS from the UKB.

**Table S3 -** Initial deep learning QC

This table contains the number of patients removed from each QC step before landmark estimation.

**Table S4 -** Image filtering

This table contains the number of patients excluded at each step of quality control following landmark estimation.

**Table S5 -** Human annotation vs model prediction

This table contains the error between human annotation and the first model prediction, as well as the error between the first model prediction and the second model prediction.

**Table S6 -** Image pixel data

This table contains the number of full-body skeletal DXA images for each pixel aspect ratio in the UKB.

**Table S7 -** Image scaling coefficient

This table contains the scaling factor estimated from the regression analysis, which is used to convert pixels to centimeters.

**Table S8 -** 7 Pelvic phenotype values across 39413 individuals

This table contains a list of all generated IDPs.

**Table S9 -** Pelvic phenotypes summary

This table contains the basic statistics of IDPs.

**Table S10 -** GCTA and LDSC heritability estimation

This table contains the heritability for each IDP as determined by GCTA.

**Table S11 -** Clumped independent SNPs and corresponding genes

This table contains output from PLINK --clump ranges command including lead SNP, p-value, the number of kilobases in each clump, gene mapping for each clump range as well as whether the single clump range genes are related to known mouse phenotypes and rare human disease.

**Table S12 -** ICD10 Codes

This table contains all ICD10 codes used in our analyses.

**Table S13 -** UKB phenotypes FID

This table contains the FID of each UKB trait used in our analyses.

**Table S14 -** Association analysis covariates

This table contains the list of covariates used in our regression analyses and the FID from the UKB

**Table S15 -** Phenotypic association results

This table contains the results from the phenotypic association analysis.

**Table S16 -** PRS association results

This table contains the results from the PRS association analysis.

**Table S17 -** Female PRS association results

This table contains the results from the female PRS association analysis.

**Table S18 -** Female genetic correlation results

This table contains the results from the female genetic correlation analysis.

**Table S19 -** MAGMA with GTEx v8

This table contains the results from the MAGMA analysis with gene expression data from GTEx v8.

**Table S20 -** MAGMA analysis across pelvic subelements across different time points

This table contains the results from the MAGMA analysis with gene expression data from different parts of the pelvis across different development time points.

**Table S21 -** MAGMA analysis across pelvic subelements

This table contains the results from the MAGMA analysis with gene expression data from different parts of the pelvis.

## Acknowledgments

This research was conducted using the UKB Resource under application no. 65439.

## Author contributions

L.X. and V.M.N. wrote the paper with input from all co-authors. L.X.,

E.K. performed analysis. L.X., E.K., D.P., and J.W. performed data preprocessing. M.B. and T.S. provided comments and helped supervise the work.

## Data and materials availability

Code used for performing the deep learning–based key point identification and quality control of the DXA data is available at https://github.com/xliaoyi/Human-pelvic-form. Our GWAS summary statistics are available at https://utexas.box.com/s/w1n8oz61sb7km2yotqyyg8em2td7b2wb. Individual-level information of skeletal lengths has been reported back to the UKB and will be available via the Access Management System.

## References

1. C. L. Lewis, N. M. Laudicina, A. Khuu, K. L. Loverro, The Human Pelvis: Variation in Structure and Function During Gait. The Anatomical Record 300, 633–642 (2017).

2. J. M. DeSilva, K. R. Rosenberg, Anatomy, Development, and Function of the Human Pelvis. The Anatomical Record 300, 628–632 (2017).

3. C. Owen Lovejoy, The natural history of human gait and posture: Part 2. Hip and thigh. Gait & Posture 21, 113–124 (2005).

4. D. M. Bramble, D. E. Lieberman, Endurance running and the evolution of Homo. Nature 432, 345–352 (2004).

5. L. Betti, Human Variation in Pelvic Shape and the Effects of Climate and Past Population History. The Anatomical Record 300, 687–697 (2017).

6. N. D. S. Grunstra, L. Betti, B. Fischer, M. Haeusler, M. Pavlicev, E. Stansfield, W. Trevathan, N. M. Webb, J. C. K. Wells, K. R. Rosenberg, P. Mitteroecker, There is an obstetrical dilemma: Misconceptions about the evolution of human childbirth and pelvic form. American Journal of Biological Anthropology 181, 535–544 (2023).

7. M. Haeusler, N. D. S. Grunstra, R. D. Martin, V. A. Krenn, C. Fornai, N. M. Webb, The obstetrical dilemma hypothesis: there’s life in the old dog yet. Biol Rev Camb Philos Soc 96, 2031–2057 (2021).

8. W. M. Krogman, The Scars of Human Evolution. Scientific American 185, 54–57 (1951).

9. M. Pavličev, R. Romero, P. Mitteroecker, Evolution of the human pelvis and obstructed labor: New explanations of an old obstetrical dilemma. Am J Obstet Gynecol 222, 3–16 (2020).

10. K. Rosenberg, W. Trevathan, Bipedalism and human birth: The obstetrical dilemma revisited. Evolutionary Anthropology: Issues, News, and Reviews 4, 161–168 (1995).

11. K. R. R. Trevathan Wenda R., “The obstetrical dilemma revisited—revisited” in The Routledge Handbook of Anthropology and Reproduction (Routledge, 2021).

12. C. M. Wall-Scheffler, H. K. Kurki, B. M. Auerbach, The Evolutionary Biology of the Human Pelvis: An Integrative Approach (Cambridge University Press, 2020).

13. S. L. Washburn, Tools and human evolution. Sci Am 203, 63–75 (1960).

14. J. C. K. Wells, J. M. DeSilva, J. T. Stock, The obstetric dilemma: An ancient game of Russian roulette, or a variable dilemma sensitive to ecology? American Journal of Physical Anthropology 149, 40–71 (2012).

15. E. Pomeroy, J. C. K. Wells, J. T. Stock, “Obstructed Labour: The Classic Obstetric Dilemma and Beyond” in Evolutionary Thinking in Medicine: From Research to Policy and Practice, A. Alvergne, C. Jenkinson, C. Faurie, Eds. (Springer International Publishing, Cham, 2016; 10.1007/978-3-319-29716-3_3), pp. 33–45.

16. J. C. K. Wells, The New “Obstetrical Dilemma”: Stunting, Obesity and the Risk of Obstructed Labour. The Anatomical Record 300, 716–731 (2017).

17. H. M. Dunsworth, There Is No “Obstetrical Dilemma”: Towards a Braver Medicine with Fewer Childbirth Interventions. Perspectives in Biology and Medicine 61, 249–263 (2018).

18. L. T. Gruss, R. Gruss, D. Schmitt, Pelvic Breadth and Locomotor Kinematics in Human Evolution. The Anatomical Record 300, 739–751 (2017).

19. A. G. Warrener, K. L. Lewton, H. Pontzer, D. E. Lieberman, A Wider Pelvis Does Not Increase Locomotor Cost in Humans, with Implications for the Evolution of Childbirth. PLOS ONE 10, e0118903 (2015).

20. C. M. Wall-Scheffler, M. J. Myers, The Biomechanical and Energetic Advantages of a Mediolaterally Wide Pelvis in Women. Anat Rec (Hoboken) 300, 764–775 (2017).

21. M. Vidal-Cordasco, A. Mateos, G. Zorrilla-Revilla, O. Prado-Nóvoa, J. Rodríguez, Energetic cost of walking in fossil hominins. American Journal of Physical Anthropology 164, 609–622 (2017).

22. P. A. Kramer, A. D. Sylvester, Hip width and metabolic energy expenditure of abductor muscles. PLOS ONE 18, e0284450 (2023).

23. J. Gorman, C. A. Roberts, S. Newsham, G. R. Bentley, Squatting, pelvic morphology and a reconsideration of childbirth difficulties. Evol Med Public Health 10, 243–255 (2022).

24. P. K. Stone, Biocultural perspectives on maternal mortality and obstetrical death from the past to the present. American Journal of Physical Anthropology 159, 150–171 (2016).

25. D. Walrath, “Bones, biases, and birth: Excavating contemporary gender norms from reproductive bodies of the past” in Exploring Sex and Gender in Bioarchaeology (2017), pp. 15–49.

26. N. D. S. Grunstra, F. E. Zachos, A. N. Herdina, B. Fischer, M. Pavličev, P. Mitteroecker, Humans as inverted bats: A comparative approach to the obstetric conundrum. American Journal of Human Biology 31, e23227 (2019).

27. J. A. Ashton-Miller, J. O. L. DeLancey, Functional anatomy of the female pelvic floor. Ann N Y Acad Sci 1101, 266–296 (2007).

28. M. M. Abitbol, Evolution of the ischial spine and of the pelvic floor in the Hominoidea. Am J Phys Anthropol 75, 53–67 (1988).

29. H. M. Dunsworth, A. G. Warrener, T. Deacon, P. T. Ellison, H. Pontzer, Metabolic hypothesis for human altriciality. Proceedings of the National Academy of Sciences 109, 15212–15216 (2012).

30. R. D. Martin, The evolution of human reproduction: A primatological perspective. American Journal of Physical Anthropology 134, 59–84 (2007).

31. K. R. Rosenberg, The evolution of modern human childbirth. American Journal of Physical Anthropology 35, 89–124 (1992).

32. H. Dunsworth, “There is no evolutionary ‘obstetrical dilemma’” in The Routledge Handbook of Anthropology and Reproduction (Routledge, 2021).

33. M. Young, D. Richard, M. Grabowski, B. M. Auerbach, B. S. de Bakker, J. Hagoort, P. Muthuirulan, V. Kharkar, H. K. Kurki, L. Betti, L. Birkenstock, K. L. Lewton, T. D. Capellini, The developmental impacts of natural selection on human pelvic morphology. Science Advances 8, eabq4884 (2022).

34. M. Young, L. Selleri, T. D. Capellini, “Chapter Nine - Genetics of scapula and pelvis development: An evolutionary perspective” in Current Topics in Developmental Biology, D. M. Wellik, Ed. (Academic Press, 2019)vol. 132 of *Organ Development*, pp. 311–349.

35. T. D. Capellini, K. Handschuh, L. Quintana, E. Ferretti, G. Di Giacomo, S. Fantini, G. Vaccari, S. L. Clarke, A. M. Wenger, G. Bejerano, J. Sharpe, V. Zappavigna, L. Selleri, Control of Pelvic Girdle Development by Genes of the Pbx Family and Emx2. Dev Dyn 240, 1173–1189 (2011).

36. E. Kun, E. M. Javan, O. Smith, F. Gulamali, J. de la Fuente, B. I. Flynn, K. Vajrala, Z. Trutner, P. Jayakumar, E. M. Tucker-Drob, M. Sohail, T. Singh, V. M. Narasimhan, The genetic architecture and evolution of the human skeletal form. Science 381, eadf8009 (2023).

37. K. Sun, Y. Zhao, B. Jiang, T. Cheng, B. Xiao, D. Liu, Y. Mu, X. Wang, W. Liu, J. Wang, High-Resolution Representations for Labeling Pixels and Regions. arXiv arXiv:1904.04514 [Preprint] (2019). 10.48550/arXiv.1904.04514.

38. J. Yang, S. H. Lee, M. E. Goddard, P. M. Visscher, GCTA: A Tool for Genome-wide Complex Trait Analysis. The American Journal of Human Genetics 88, 76–82 (2011).

39. B. Fischer, P. Mitteroecker, Allometry and Sexual Dimorphism in the Human Pelvis. The Anatomical Record 300, 698–705 (2017).

40. S. L. Cox, A geometric morphometric assessment of shape variation in adult pelvic morphology. American Journal of Physical Anthropology 176, 652–671 (2021).

41. B. Fischer, P. Mitteroecker, Covariation between human pelvis shape, stature, and head size alleviates the obstetric dilemma. Proceedings of the National Academy of Sciences 112, 5655–5660 (2015).

42. P.-R. Loh, G. Tucker, B. K. Bulik-Sullivan, B. J. Vilhjálmsson, H. K. Finucane, R. M. Salem, D. I. Chasman, P. M. Ridker, B. M. Neale, B. Berger, N. Patterson, A. L. Price, Efficient Bayesian mixed-model analysis increases association power in large cohorts. Nat Genet 47, 284–290 (2015).

43. Z. R. McCaw, R. Dey, H. Somineni, D. Amar, S. Mukherjee, K. Sandor, insitro Research Team, T. Karaletsos, D. Koller, G. Davey Smith, D. MacArthur, C. O’Dushlaine, T. W. Soare, Pitfalls in performing genome-wide association studies on ratio traits. [Preprint] (2023). 10.1101/2023.10.27.564385.

44. B. K. Bulik-Sullivan, P.-R. Loh, H. K. Finucane, S. Ripke, J. Yang, N. Patterson, M. J. Daly, A. L. Price, B. M. Neale, LD Score regression distinguishes confounding from polygenicity in genome-wide association studies. Nat Genet 47, 291–295 (2015).

45. E. Lausch, P. Hermanns, H. F. Farin, Y. Alanay, S. Unger, S. Nikkel, C. Steinwender, G. Scherer, J. Spranger, B. Zabel, A. Kispert, A. Superti-Furga, *TBX15* Mutations Cause Craniofacial Dysmorphism, Hypoplasia of Scapula and Pelvis, and Short Stature in Cousin Syndrome. The American Journal of Human Genetics 83, 649–655 (2008).

46. E. M. H. F. Bongers, P. H. G. Duijf, S. E. M. Van Beersum, J. Schoots, A. Van Kampen, A. Burckhardt, B. C. J. Hamel, F. Lošan, L. H. Hoefsloot, H. G. Yntema, N. V. A. M. Knoers, H. Van Bokhoven, Mutations in the Human TBX4 Gene Cause Small Patella Syndrome. The American Journal of Human Genetics 74, 1239–1248 (2004).

47. T. Ge, C.-Y. Chen, Y. Ni, Y.-C. A. Feng, J. W. Smoller, Polygenic prediction via Bayesian regression and continuous shrinkage priors. Nat Commun 10, 1776 (2019).

48. Y. Melesse, T. Assebe Yadeta, M. Lami, T. Getachew, H. Mohammed, B. Berhanu, M. Dheresa, One-sixth of women experienced obstructed labor among those delivered at public hospitals in Southern Ethiopia: A multicenter study. SAGE Open Med 11, 20503121231164056 (2023).

49. M. Desta, Z. Mekonen, A. A. Alemu, M. Demelash, T. Getaneh, Y. Bazezew, G. M. Kassa, N. Wakgari, Determinants of obstructed labour and its adverse outcomes among women who gave birth in Hawassa University referral Hospital: A case-control study. PLoS One 17, e0268938 (2022).

50. A. Portmann, Die Tragzeiten Der Primaten Und Die Dauer Der Schwangerschaft Beim Menschen: Ein Problem Der Vergleichenden Biologie (Verlag nicht ermittelbar, 1941).

51. S. J. Gould, Ever Since Darwin: Reflections in Natural History (W. W. Norton & Company, 1992).

52. E. Pomeroy, J. T. Stock, T. J. Cole, M. O’Callaghan, J. C. K. Wells, Relationships between Neonatal Weight, Limb Lengths, Skinfold Thicknesses, Body Breadths and Circumferences in an Australian Cohort. PLoS One 9, e105108 (2014).

53. J. G. Alves, L. C. Siqueira, L. M. Melo, J. N. Figueiroa, Smaller pelvic size in pregnant adolescents contributes to lower birth weight. International Journal of Adolescent Medicine and Health 25, 139–142 (2013).

54. I. R. Timmins, F. Zaccardi, C. P. Nelson, P. W. Franks, T. Yates, F. Dudbridge, Genome-wide association study of self-reported walking pace suggests beneficial effects of brisk walking on health and survival. Commun Biol 3, 1–9 (2020).

55. K. Norman, N. Stobäus, M. C. Gonzalez, J.-D. Schulzke, M. Pirlich, Hand grip strength: Outcome predictor and marker of nutritional status. Clinical Nutrition 30, 135–142 (2011).

56. F. Zaccardi, I. R. Timmins, J. Goldney, F. Dudbridge, P. C. Dempsey, M. J. Davies, K. Khunti, T. Yates, Self-reported walking pace, polygenic risk scores and risk of coronary artery disease in UK biobank. Nutrition, Metabolism and Cardiovascular Diseases 32, 2630–2637 (2022).

57. C. A. Celis-Morales, S. Gray, F. Petermann, S. Iliodromiti, P. Welsh, D. M. Lyall, J. Anderson, P. Pellicori, D. F. Mackay, J. P. Pell, N. Sattar, J. M. R. Gill, Walking Pace Is Associated with Lower Risk of All-Cause and Cause-Specific Mortality. Medicine & Science in Sports & Exercise 51, 472 (2019).

58. A. M. Alaa, T. Bolton, E. Di Angelantonio, J. H. F. Rudd, M. Van Der Schaar, Cardiovascular disease risk prediction using automated machine learning: A prospective study of 423,604 UK Biobank participants. PLoS ONE 14, e0213653 (2019).

59. J. Boonpor, S. Parra-Soto, J. Gore, A. Talebi, N. Lynskey, A. Raisi, P. Welsh, N. Sattar, J. P. Pell, J. M. R. Gill, S. R. Gray, F. K. Ho, C. A. Celis-Morales, Association between walking pace and incident type 2 diabetes by adiposity level: A prospective cohort study from the UK Biobank. Diabetes, Obesity and Metabolism 25, 1900–1910 (2023).

60. H. E. Syddall, L. D. Westbury, C. Cooper, A. A. Sayer, Self-reported walking speed: a useful marker of physical performance among community-dwelling older people? J Am Med Dir Assoc 16, 323–328 (2015).

61. A. Zeki Al Hazzouri, E. R. Mayeda, T. Elfassy, A. Lee, M. C. Odden, D. Thekkethala, C. B. Wright, M. M. Glymour, M. N. Haan, Perceived Walking Speed, Measured Tandem Walk, Incident Stroke, and Mortality in Older Latino Adults: A Prospective Cohort Study. The Journals of Gerontology: Series A 72, 676–682 (2017).

62. A. V. Rowlands, P. C. Dempsey, B. Maylor, C. Razieh, F. Zaccardi, M. J. Davies, K. Khunti, T. Yates, Self-reported walking pace: A simple screening tool with lowest risk of all-cause mortality in those that “walk the talk.” J Sports Sci 41, 333–341 (2023).

63. A. Huseynov, C. P. E. Zollikofer, W. Coudyzer, D. Gascho, C. Kellenberger, R. Hinzpeter, M. S. Ponce de León, Developmental evidence for obstetric adaptation of the human female pelvis. Proceedings of the National Academy of Sciences 113, 5227–5232 (2016).

64. C. Bycroft, C. Freeman, D. Petkova, G. Band, L. T. Elliott, K. Sharp, A. Motyer, D. Vukcevic, O. Delaneau, J. O’Connell, A. Cortes, S. Welsh, A. Young, M. Effingham, G. McVean, S. Leslie, N. Allen, P. Donnelly, J. Marchini, The UK Biobank resource with deep phenotyping and genomic data. Nature 562, 203–209 (2018).

65. C. R. Harris, K. J. Millman, S. J. van der Walt, R. Gommers, P. Virtanen, D. Cournapeau, E. Wieser, J. Taylor, S. Berg, N. J. Smith, R. Kern, M. Picus, S. Hoyer, M. H. van Kerkwijk, M. Brett, A. Haldane, J. F. del Río, M. Wiebe, P. Peterson, P. Gérard-Marchant, K. Sheppard, T. Reddy, W. Weckesser, H. Abbasi, C. Gohlke, T. E. Oliphant, Array programming with NumPy. Nature 585, 357–362 (2020).

66. P. Virtanen, R. Gommers, T. E. Oliphant, M. Haberland, T. Reddy, D. Cournapeau, E. Burovski, P. Peterson, W. Weckesser, J. Bright, S. J. van der Walt, M. Brett, J. Wilson, K. J. Millman, N. Mayorov, A. R. J. Nelson, E. Jones, R. Kern, E. Larson, C. J. Carey, İ. Polat, Y. Feng, E. W. Moore, J. VanderPlas, D. Laxalde, J. Perktold, R. Cimrman, I. Henriksen, E. A. Quintero, C. R. Harris, A. M. Archibald, A. H. Ribeiro, F. Pedregosa, P. van Mulbregt, SciPy 1.0: fundamental algorithms for scientific computing in Python. Nat Methods 17, 261–272 (2020).

67. S. van der Walt, J. L. Schönberger, J. Nunez-Iglesias, F. Boulogne, J. D. Warner, N. Yager, E. Gouillart, T. Yu, scikit-image: image processing in Python. PeerJ 2, e453 (2014).

68. T.-Y. Lin, M. Maire, S. Belongie, L. Bourdev, R. Girshick, J. Hays, P. Perona, D. Ramanan, C. L. Zitnick, P. Dollár, Microsoft COCO: Common Objects in Context. arXiv arXiv:1405.0312 [Preprint] (2015). 10.48550/arXiv.1405.0312.

69. K. Sun, B. Xiao, D. Liu, J. Wang, Deep High-Resolution Representation Learning for Human Pose Estimation. arXiv arXiv:1902.09212 [Preprint] (2019). 10.48550/arXiv.1902.09212.

70. C. C. Chang, C. C. Chow, L. C. Tellier, S. Vattikuti, S. M. Purcell, J. J. Lee, Second-generation PLINK: rising to the challenge of larger and richer datasets. GigaScience 4, s13742–015-0047–8 (2015).

71. D. M. Church, V. A. Schneider, T. Graves, K. Auger, F. Cunningham, N. Bouk, H.-C. Chen, R. Agarwala, W. M. McLaren, G. R. S. Ritchie, D. Albracht, M. Kremitzki, S. Rock, H. Kotkiewicz, C. Kremitzki, A. Wollam, L. Trani, L. Fulton, R. Fulton, L. Matthews, S. Whitehead, W. Chow, J. Torrance, M. Dunn, G. Harden, G. Threadgold, J. Wood, J. Collins, P. Heath, G. Griffiths, S. Pelan, D. Grafham, E. E. Eichler, G. Weinstock, E. R. Mardis, R. K. Wilson, K. Howe, P. Flicek, T. Hubbard, Modernizing Reference Genome Assemblies. PLOS Biology 9, e1001091 (2011).

72. S. Purcell, B. Neale, K. Todd-Brown, L. Thomas, M. A. R. Ferreira, D. Bender, J. Maller, P. Sklar, P. I. W. de Bakker, M. J. Daly, P. C. Sham, PLINK: A Tool Set for Whole-Genome Association and Population-Based Linkage Analyses. The American Journal of Human Genetics 81, 559–575 (2007).

73. K. Watanabe, E. Taskesen, A. van Bochoven, D. Posthuma, Functional mapping and annotation of genetic associations with FUMA. Nat Commun 8, 1826 (2017).

74. C. Palazzo, C. Nguyen, M.-M. Lefevre-Colau, F. Rannou, S. Poiraudeau, Risk factors and burden of osteoarthritis. Annals of Physical and Rehabilitation Medicine 59, 134–138 (2016).

